# Informational Complexity as a Neural Marker of Cognitive Reserve

**DOI:** 10.1101/2025.02.13.638084

**Authors:** Laura Stolp, Kanad N Mandke, Pedro AM Mediano, Helena M Gellersen, Alex Swartz, Katarzyna Rudzka, Jon Simons, Tristan A Bekinschtein, Daniel Bor

## Abstract

In Alzheimer’s disease (AD), a mismatch between neurological damage and cognitive functioning often is attributed to individual differences in cognitive reserve. Understanding the neural mechanism of cognitive reserve could help assessing the therapeutic effectiveness of interventions in AD. To address this, here, 38 elderly participants performed a sustained attention task during high-density EEG while awake and during drowsiness. Operationally, the degree to which performance was impaired under drowsiness signalled the extent of cognitive reserve, with less impairment indicating a higher level of cognitive reserve. Investigating performance variations during the active management of neural challenges offers a novel approach to studying cognitive reserve, capturing dynamics that mirror everyday cognitive demand. We related cognitive reserve to various measures, including informational complexity using the Lempel-Ziv (LZSUM) algorithm. We found a significant interaction effect between arousal and performance, where LZSUM values increased in high performers when drowsy but decreased in low performers. This effect was most pronounced in the frontal and central areas. Our findings suggest LZSUM to be indicative of a compensatory mechanism and thus show potential for LZSUM as a neural marker in assessing cognitive reserve. However, we found no consistent relationship between performance and structural brain measures, and proxies of cognitive reserve. Critically, our findings present a counterexample to the prevailing view that informational complexity purely reflects conscious level. Further research, such as a study with the same paradigm in patients with mild cognitive impairment (MCI) and AD, may lead to additional insights of whether we are truly measuring cognitive reserve.

## 1. INTRODUCTION

Dementia is a neurodegenerative disease that causes a decline in cognitive function. Patients experience deficits in memory, language abilities, learning capacity, and executive processing. These impairments result in significant difficulties in performing everyday activities (Duthey, 2013). Alzheimer’s disease (AD) is the most common form of dementia and accounts for up to 70% of dementia cases (WHO, 2019). Interestingly, there is not always a direct relationship between the level of cognitive decline and neuropathological severity (Katzman et al., 1988). This phenomenon may be explained by a compensatory mechanism known as “cognitive reserve” (Stern, 2012). Although cognitive reserve is most frequently applied to dementia, the concept of cognitive reserve aims to explain individual variability in cognitive functioning and clinical status in the broader context of any neurological or psychiatric strain. In high cognitive reserve dementia patients, advanced neurodegeneration has been observed in concurrence with relatively well-preserved cognition (Steffener & Stern, 2012).

In recent years, most AD clinical trials have failed to show significant differences between drug and placebo groups, a situation that may be explained by the irreversible neurological damage that may be present many years before AD symptoms emerge (Cummings et al., 2018). Disease progression markedly differs between patients with high and low cognitive reserve. In patients with high cognitive reserve clinical symptoms will be presenting much later in the disease progression compared to low cognitive reserve patients, and thus, they can maintain their cognitive functioning for longer. However, once symptoms appear, their cognitive decline accelerates and is often more rapid than in patients with low cognitive reserve (Stern, 2012). The masking effect of high cognitive reserve on the severity of the condition necessitates more precise diagnostic methods for these patients. Furthermore, when the disease is caught at an earlier stage, treatment with currently available medication may be more successful and prevent further damage. For this very reason, current research into pharmacotherapy for AD is focused on the preclinical stage where pathology is present in the absence of neuropsychological symptoms (Huang et al., 2023). In clinical trials, it is crucial to consider and stratify cognitive reserve, since these trials often assess differences in the rate of decline in patients to determine drug treatment effectiveness compared to a placebo. If the level of cognitive reserve is not accounted for, cognitive reserve could be a strong confounding factor in clinical research, which might lead to incorrect conclusions about the (in)effectiveness of a given treatment. Therefore, a direct measure to assess a patient’s level of cognitive reserve would be beneficial.

Closely related to cognitive reserve is the concept of brain reserve. Brain reserve refers to structural reserve in the form of higher brain volume and greater neural density. Individuals with this type of surplus capacity are better able to cope with the strain age- or disease related changes and maintain cognitive functioning for longer (Fratiglioni & Wang, 2007). In contrast to the *passive* construct of brain reserve, *cognitive reserve* is defined in terms of *active* adaptability (i.e. flexibility, capacity, efficiency) of functional brain networks and neural communication. With high adaptability the brain actively resists the effects of ageing and pathology, and preserves cognitive functioning (Stern et al., 2020). Just as individuals differ in their structural reserve of the brain, there are individual differences in the adaptability of functional brain processes. Traditionally, cognitive reserve has been associated with various demographic and lifestyle characteristics (Satz, 1993; Scarmeas et al., 2001; Valenzuela et al., 2008). Proxies of cognitive reserve such as occupational complexity, educational level or premorbid IQ have been shown to moderate the relationship between neural changes and clinical status but are not direct measures of cognitive reserve. It is not clear whether and how these factors are fundamental to the construct of cognitive reserve. As a result of the uncertainty about its precise nature, cognitive reserve has often been measured as a latent variable (Jones et al., 2011; Stern et al., 2019).

A way to measure cognitive reserve more directly may be to temporarily strain the neural system and compare baseline task performance against performance under neural strain. For instance, inducing drowsiness experimentally could be a suitable approach. Drowsiness is a reversible and common way of straining the neural system (Nilsson et al., 2005; Goupil & Bekinschtein, 2012). High levels of drowsiness are associated with cognitive impairment (Durmer & Dinges, 2005; Lacaux et al., 2024). The degree to which an individual experiences cognitive impairment due to drowsiness may reflect individual differences in cognitive reserve. Moreover, AD symptoms and sleep disturbances are highly associated factors. Furthermore, sleep problems are one of the first emerging symptoms of AD, suggesting that brain areas related to circadian control and sleep are impacted early in the disease pathogenesis. Individuals suffering from mild cognitive impairment, often an early sign of AD, exhibit certain EEG abnormalities, such as a reduction in slow-wave-sleep (SWS) and sleep spindles (Ju et al., 2014). Sleep disturbances can lead to excessive daytime napping, drowsiness, and an increase in cognitive symptoms (Moran et al., 2005; Ju et al, 2014). The importance of sleep disturbances in AD, coupled with the notion that the degree of drowsiness-induced cognitive impairment might reflect variations in cognitive reserve, suggests that level of arousal could be a useful experimental manipulation in the context of AD research and cognitive reserve.

Many studies have investigated how cognitive reserve is implemented on a neural level (Stern et al., 2019). As our understanding advances, the aim is to transition from proxy measures to direct neural markers that may aid a more accurate assessment of cognitive reserve. Since cognitive reserve is defined in terms of adaptability (i.e. flexibility, capacity, efficiency) of brain networks and neural communication (Stern et al., 2019), high cognitive reserve might be related to generally more efficient information processing. A novel line of research involves quantifying informational complexity in the brain and relating this to different levels of (un)consciousness (Schartner et al., 2015; Chennu et al., 2014; Casali et al., 2013; Pascovich et al., 2022). Furthermore, recent research indicates individual differences in performance under neural strain are related to informational complexity (Boncompte et al., 2021). In a study that examined informational complexity of electroencephalograph (EEG) signals when participants underwent mild propofol sedation while performing an auditory discrimination task, high performers were shown to maintain or increase informational complexity, whereas informational complexity decreased in low performers. This effect was most pronounced in the frontal regions (Boncompte et al., 2021), which is noteworthy given the involvement of the frontal regions in attention, executive processes and cognitive control (Kievit et al., 2014; Stern et al., 2019). Given these findings, informational complexity could be a promising neural correlate for assessing cognitive reserve. A well-established approach to measuring informational complexity is by using the Lempel-Ziv (LZUM) algorithm, which assesses the compressibility of a given signal such as an EEG recording. Higher compressibility of a signal indicates lower informational complexity and vice versa (Lempel & Ziv, 1976). LZSUM has been used extensively in consciousness studies and has more recently been applied to research on individual differences in informational complexity under neural strain in healthy young adult participants (Boncompte et al., 2021).

The current study aimed to measure cognitive reserve directly and relate it to a range of neural measures. In this EEG study, 38 elderly healthy participants who already had undergone structural MRI scans carried out a sustained attention task under two arousal states: alert and drowsy. Task performance metrics included mean reaction time, variation of reaction time, and error-based metrics. Here, the level of cognitive reserve was operationally defined by assessing to what degree task performance was impaired under drowsiness, compared to alertness. We hypothesised LZSUM values to decrease under drowsiness in low performers but expected similar or even higher complexity values in high performers when comparing drowsiness to alertness, especially in frontal regions, in line with previous research (Boncompte et al., 2021; Kievit et at., 2014). Additionally, we examined the relationship between cognitive reserve and both structural volume in cortical and hippocampal areas, and network analyses of white matter connectivity. This approach aims to provide a comprehensive understanding of the potential neural underpinnings of cognitive reserve, so we can draw more robust conclusions from a variety of neuroimaging metrics. By using performance under neural strain (i.e. drowsiness) to define cognitive reserve and using informational complexity measures, this study aims to fill a critical gap in the literature by introducing a more direct measure of cognitive reserve, grounded in neurophysiology. Finally, incorporating both functional and structural neural markers provides a comprehensive exploration of cognitive reserve, which could aid in the development of targeted therapeutic interventions in Alzheimer’s disease and related dementias.

## 2. METHODS

### 2.1 Participants

38 elderly participants (16 females, 22 males) with an average age of 73.16 years (SD = 5.26, range 60-84) performed the auditory version of the Sustained Attention to Response Task (SART) (Seli et al., 2012) while their EEG was recorded. For this study, we recruited participants whose structural magnetic resonance imaging (MRI) scans (T1-weighted and diffusion weighted imaging) had already been obtained through collaborative research projects conducted within two years before the data collection of the present study (Gellersen et al., 2023). The experimental procedure took place at the EEG lab of the Consciousness and Cognition research group of the University of Cambridge. On session days, participants were asked to refrain from the consumption of any caffeinated beverages. Participants were paid £10 per hour as reimbursement for their time. The experiment lasted approximately 3.5 hours in total including EEG setup, experiment, and cognitive tests. Before the experiment, participants gave their written informed consent. Ethical approval was given by the Psychological Research Ethics Committee at the University of Cambridge (CPREC2014.25).

Before the start of the experimental task, the participants were subjected to a battery of cognitive tests. Fluid intelligence was determined by participant performance on the Cattell Culture Fair Intelligence Test (scale 2A, Cattell, 1973). Premorbid IQ (i.e. WAIS IQ) was estimated using the National Adult Reading Test (NART). (Nelson & Willison, 1991). Additionally, we included participants’ scores from the Montreal Cognitive Assessment (MoCA), which had been conducted two years prior during the initial study visit (Gellersen et al., 2023). The MoCA is a brief 10-minute screening tool to assess cognitive functioning with a maximum score of 30 points. It is often used by first-line clinicians in the assessment of patients with mild cognitive complaints. The test consists of several parts to assess various cognitive domains, including memory, executive functioning, attention, and language. (Nasreddine et al., 2005). A score of 26 points or above is considered normal, while a score lower than 26 indicates cognitive impairment, with lower scores representing worse impairment. In our cohort, 6 participants scored below 26 points. Furthermore, on average there was a stark difference between current fluid IQ (Cattell) and NART IQ, with the latter being considerable higher. Since healthy ageing is a cause of neural strain (Whalley et al., 2004), the observed difference in our participant group between estimated premorbid IQ and current levels of cognitive functioning aligns with expectations. See Table 1 for a detailed summary of participant characteristics.

**Table 1.** Participant Characteristics.

| **N = 38** (♀ = 16, ♂ = 22) | **Mean (SD)** | **Median** | **Range** |
| --- | --- | --- | --- |
| Age | 73.16 (5.26) | 73.5 | 60 – 84 |
| Years of Education | 18.01 (4.68) | 18 | 10 – 39 |
| MoCA score | 27 (1.76) | 27 | 23 – 30 |
| NART IQ | 120.63 (6.02) | 121 | 105 – 129 |
| Cattell IQ | 97.18 (10.49) | 96 | 79 – 122 |
| Difference NART – Cattell | 23.48 (11.74) | 25.5 | -10 – 42 |

### 2.2 Experimental procedure

In the auditory version of the SART, participants were instructed to respond to certain auditory stimuli and inhibit responses in reaction to others (Seli et al., 2012). As part of the experimental setup, levels of arousal were manipulated in the following way. Before the task, participants were seated in a reclining chair in a darkened room and provided with blankets and pillows to ensure their comfort. The participants were given the following instructions: a) to close their eyes and keep them closed throughout the task, b) to minimise any movements, c) to continue the task even if they made mistakes, and, d) not to worry if they felt drowsy during the task. However, if a participant fell asleep, as indicated by three consecutive non-responsive trials, an audio recording was played to wake them up. If the recording failed to wake the participant, the experimenter manually woke them up. The purpose of this setup was to induce drowsiness. The expectation was that participants would be relatively alert at the start of the task, but increasingly drowsy as the experiment progressed, aiding the comparison of performance in alert and drowsy states.

Prior to the onset of the SART, participants completed a resting state block (5 min). The SART involved the randomised auditory presentation of numbers between 0 and 9, extracted from a vocal recording database (Sayanng, 2009). Participants were instructed to respond to each number by pressing a button, except for the number ‘3’, the target stimulus, to which they had to withhold a response. Stimulus randomisation was modulated by two parameters: first, the target stimulus was not presented more than three times consecutively; second, the target stimulus was only presented in 10% of trials. Participants initially performed a practice block consisting of 30 trials (2.5 min), with a fixed response window of 1100ms. For the main task, the duration of the response window was determined by the mean reaction time of the participants during the practice block (i.e. mean RT + 250 ms). The end of the response window was signalled by a ‘beep’ sound (SoundJay, 2019), followed by an inter-trial interval lasting between 2 and 5 seconds. The task consisted of 700 trials in total and was divided into two blocks of 350 trials each, with a 3-minute break in between.

### 2.3 EEG DETAILS

#### 2.3.1 EEG acquisition details

A Philips EGI EEG system with 129 channels (Electrical Geodesics, Inc.) was used to collect EEG data. The recordings were obtained with Netstation software running on a Mac computer and sampled at a frequency of 1000Hz.

#### 2.3.2 EEG preprocessing

Down-sampling of the data to 250 Hz was done using Netstation Tools software (EGI, Electrical Geodesics, Inc.) after which the data was imported to EEGLAB (Delorme and Makeig, 2004) for MATLAB format (version 2019a, MathWorks, Inc) using the mffmatlabio plugin (Delorme et al., 2019). The Automagic system (Pedroni, Bahreini, and Langer, 2019) was used to standardise the pre-processing pipeline. First, the PREP algorithm (Bigdely-Shamlo et al., 2015) was used to detect and remove noisy channels. The remaining channels were then filtered using a high-pass cut-off of 1.0 Hz and a low-pass cut-off of 35 Hz. Next, an independent components analysis was performed on the data. Artefactual components were automatically rejected using the open-source EEGLAB plug-in MARA (Winkler et al., 2014). Removed channels (mean = 11.42, SD = 7.25) were then spherically interpolated. The data was pre-trial epoched into periods of four seconds preceding each stimulus onset, and noisy epochs were automatically rejected (mean = 6, SD = 4.31) based on threshold values of ±150 μV or slopes of more than 60 μV using the manage badTrials plugin for EEGLAB (Jagannathan et al., 2018).

#### 2.3.3 Drowsiness classification

To determine the level of arousal for each 4-second epoch, a machine learning algorithm was used that had previously been validated for classifying levels of arousal (Jagannathan et al., 2018). The algorithm was trained using microstate variations in alertness and drowsiness derived from the EEG signal of eyes-closed experiments. Specifically, the algorithm is based on the Hori-scale, which divides the sleep onset process into nine stages ranging from wakefulness (stages 1-2) to the onset of N2 sleep (stage 9) (Iber et al., 2007). The algorithm computes predictor frequency variance and cross-frequency coherence features, which are then used to differentiate between alertness and drowsiness. Alertness is determined by the presence of Alpha wave trains (> 50%) (Hori: 1 – 2), whereas the transition from alertness to drowsiness is typically characterised by a reduction in Alpha activity (Hori: 3 = Alpha < 50%) and an increase in Theta waves (Hori: 5). The EEG data is then further analysed to detect graphoelements (i.e. vertex, spindles, and K-complexes) which are used to classify the level of drowsiness as either ‘mild’ when no graphoelements are present or ‘severe’ in case graphoelements do occur (Jagannathan et al., 2018). In the present study, trials classified as severely drowsy were merged with those classified as mildly drowsy due to the lack of severely drowsy trials in most participants. The resulting levels of arousal were categorised as ‘alert’ and ‘drowsy’ for analysis of within- and between-participant effects.

#### 2.3.4 Lempel-Ziv (LZ) complexity algorithm

The LZ algorithm computes complexity by determining the number of non-redundant patterns in the given data and is therefore inherently a measure of signal diversity (Lempel & Ziv, 1976). LZ is a widely recognized measure of informational complexity and has previously been applied to the EEG signal (Casali et al., 2013; Schartner et al., 2015). In our study, we applied the LZ algorithm in its single-channel form, known as LZSUM, which focuses on capturing the temporal diversity within individual EEG channels. To compute LZSUM, EEG data was first transformed into a binary sequence based on a threshold derived from the instantaneous amplitude of the Hilbert transform, following the approach used by Schartner et al. (2015). This binary representation assesses signal complexity by identifying unique patterns within the sequence. The LZSUM value, ranging from 0 (indicating no diversity) to 1 (indicating maximum diversity), provides an insight into the complexity of the EEG signal, where higher values denote a less compressible, more diverse signal. See Schartner et al. (2015) for a detailed explanation of the LZ algorithm and its application to EEG data.

### 2.4 MRI DETAILS

#### 2.4.1 MRI image acquisition

Structural MRI and Diffusion Weighted Images (DWI) were acquired at the MRC Cognition and Brain Sciences Unit of the University of Cambridge. Here, whole-brain T1-weighted (1×1×1 mm) MRI image acquisition was performed using a 3-Tesla Trio Siemens scanner (32-channel coil). As previously described in Gellersen et al. (2023, [p. 92]), the following acquisition parameters were used: “a whole-brain T1-weighted (1×1×1 mm) magnetization-prepared rapid gradient-echo (MPRAGE) sequence with a repetition time (TR) of 2300 ms, an echo time (TE) of 2.96 ms, a field of view (FOV) of 256 mm, flip angle of 9°, and 176 sagittal slices.” The acquisition was done in an interleaved, bottom-up order. Diffusion Weighted Imaging (DWI) was performed using isotropic voxel resolution (2x2x2mm3) and an interleaved slice acquisition method to minimise cross-talk artefacts between slices. The following acquisition parameters were used: a total acquisition time of 10 min and 14 sec, a repetition time (TR) of 8500 ms, an echo time (TE) of 90 ms, a field of view (FOV) of 192 mm x 192 mm, with a matrix size of 96 x 96, resulting in a resolution of 2 mm x 2 mm. Imaging was conducted in 2D with a total of 68 slices, each 2 mm thick, and without any slice gap. The sequence included 64 diffusion directions with two diffusion weightings, characterized by b-values of 0 s/mm^2^ and 1000 s/mm^2^. Parallel imaging was employed using the GRAPPA technique, with an acceleration factor in the phase-encode (PE) direction of 2 and 40 reference lines in the PE (Gellersen et al., 2023).

#### 2.4.2 Structural MRI preprocessing

In our study, structural MRI data underwent processing and parcellation to analyse cortical volumes and hippocampal subfields. We used FreeSurfer software for the segmentation of T1-weighted scans, obtaining total intracranial volume (TIV) and volumetric measures of cortical regions. The cortical regions were parcellated according to the Desikan-Killiany atlas (Desikan et al., 2006; Fischl and Dale, 2000). For more information on the cortical parcellation process, readers are referred to the FreeSurfer wiki page on Cortical Parcellation [https://surfer.nmr.mgh.harvard.edu/fswiki/CorticalParcellation]. Manual segmentations of the T2 images were previously obtained by two independent raters (authors HMG and BFG in Gellersen et al., 2023) to delineate MTL sub-regions using the ITK-Snap software (Version 3.6.0; www.itksnap.org) (Yushkevich et al., 2006) and following a protocol developed by Carr and colleagues (2017). This method allowed a precise delineation of the perirhinal cortex (PRC), entorhinal cortex (ERC), parahippocampal cortex (PHC), and the hippocampal subfields comprising the subiculum, CA1, a combined CA2-4/dentate gyrus region, and the hippocampal tail. For more details see Gellersen et al. (2023).

#### 2.4.3 Diffusion image preprocessing

For the DWI preprocessing, we used the same approach described in Luppi et al. (2021). Here, MRtrix3 tools were used to preprocess the diffusion-weighted images (Tournier et al., 2019). First, we manually removed the diffusion-weighted volumes that showed a considerable degree of distortion. After completion of this step, the pipeline consisted of the following steps: (1) Using the *dwidenoise* command, diffusion data were denoised with a technique that utilises DWI data redundancy in the PCA domain (Veraart et al., 2016), (2) The images were corrected for eddy current distortion and motion by registration of all diffusion images to b0 with the use of FSL’s *eddy* tool through the *dwipreproc* command of MRtrix3, (3) The diffusion gradient vectors were rotated to correct participant motion as estimated by the *eddy* tool, (4) Using the *dwibiascorrect* command, DWI volumes were corrected for b1 field inhomogeneity, (5) We used a combination of FSL *BET* commands and MRtrix3 *dwi2mask* to create a brain mask.

We used DSI Studio (www.dsistudio.labsolver.org) to reconstruct the diffusion tensor imaging (DTI) data from the preprocessed diffusion data. Specifically, we applied q-space diffeomorphic reconstruction (QSDR) (Yeh et al., 2011) which is a previously-validated technique to reconstruct structural networks. This model-free technique maintains the fiber geometry continuity for fiber tracking and conserves the diffusible spins by computing the orientational distribution of water diffusion density in standard space. The first step consists of the reconstruction of DWIs in native space and quantitative anisotropy (QA) calculation per voxel. Using the acquired QA values, the brains are then warped in Montreal Neurological Institute (MNI) space to a template QA volume with the nonlinear registration algorithm of the statistical parametric mapping (SPM) application. After converting the images to MNI space, the final step of the QSDR consists of reconstructing spin density functions (SDFs) using the following parameters: mean diffusion distance = 1.25mm, 3 fiber orientations for each voxel (Yeh et al., 2011).

To identify the connectivity between brain areas, we employed deterministic fiber tracking using a “FACT” algorithm with 1,000,000 streamlines. For this aim, we used the following previously-validated parameters (Luppi & Stamatakis, 2020; Medaglia, 2016) as previously described in (Luppi & Stamatakis, 2020, [p.99]) and (Luppi et al., 2021, [p. 35, *biorxiv*]): “angular cutoff = 55°, step size = 1.0 mm, tract length between 10mm (minimum) and 400mm (maximum), no spin density function smoothing, and QA threshold determined by DWI signal in the cerebrospinal fluid.” To generate a white matter mask, a standard threshold of 0.6 Otsu is automatically employed by DSI Studio to threshold the spin density function’s anisotropy values. The mask is then used to automatically check every streamline, to exclude streamlines with incorrect termination locations (Luppi & Stamatakis, 2020; Medaglia, 2016).

#### 2.4.4 Parcellation & network modules

We used a parcellation method, developed by Schaefer et al. (2018), to divide the structural MRI brains into 100 cortical regions of interest (i.e. network nodes). Next, all network nodes were allocated to one of the seven subnetworks as previously defined by Yeo et al. (2011). The different subnetworks were numbered in the following way: 1. Visual areas (VIS), 2. Somatomotor system (SOM), 3. Dorsal Attention Network (DOR), 4. Ventral Attention Network (VEN), 5. Limbic regions (LIM), 6. Frontoparietal Control Network (FPC), and 7. Default Mode Network (DMN).

### 2.5 STATISTICAL ANALYSIS

#### 2.5.1 Behavioural performance and exclusion

Statistical analyses were performed using MATLAB (version 2019b). To determine performance on the SART, we analysed the mean reaction times (RT), the coefficient of variation of reaction time (CV RT), and the proportion of commission errors and omission errors of each participant. The CV RT determined RT variability and was computed as the standard deviation divided by the mean. The proportion of commission errors was defined as the sum of all target trials with a response (i.e. errors due to responding when the participants should not have) divided by the sum of all target trials. Proportion of omission errors was defined as the sum of all non-target trials without a response (i.e. errors due to not sresponding when the participants should have) or a response made after the end of the response window, divided by the sum of all non-target trials. Target trials were excluded from further analysis when they immediately followed three succeeding non-target trials where no response was given, to ensure that seemingly correct response omissions on target trials were not due to the participant being asleep. The following exclusion criteria were used for participants: (1) Participants were excluded from all analyses involving reaction time data when they had fewer than 50 alert or drowsy trials, (2) Participants were excluded from all analyses involving commission error data when they had fewer than 10 alert or drowsy *target* trials. As a result, 34 participants were included in analyses concerning measures of RT. Similarly, for analyses concerning errors (i.e. omission and commission errors), participants with fewer than 10 *target* trials in drowsy or alert conditions were excluded leaving 28 participants included in error-related analyses. Additionally, we assessed the presence of trains of omission errors, defined as sequences of at least three consecutive omission errors, indicative of the participant being asleep rather than genuinely failing to respond. This analysis aimed to distinguish between true omission errors and those resulting from sleep. Two participants exhibited a significantly higher incidence of trains of omission errors, indicating either sleep or other forms of non-compliance, and thus were additionally excluded from analyses concerning omission errors.

#### 2.5.2 Ranking

To divide the participants into high and low performers, we calculated the difference between alert and drowsy states for each of the four-performance metrics: mean RT (drowsy/alert ratio), CV RT (drowsy/alert ratio), commission (drowsy minus alert) and omission errors (drowsy minus alert). Generally, participants performed worse while drowsy, although there were some exceptions in which the score improved while drowsy. Participants were ranked based on the performance difference between alertness and drowsiness, ranging from participants exhibiting little performance decline or even improvement under drowsiness (high performers), to those who experienced the greatest decline in performance between drowsiness and alertness (low performers). For grouping based on performance, we excluded the middle third of participants, leading to our final groupings which included 23 participants (11 high, 12 low) for mean RT and CV RT, 19 (9 high, 10 low) for commission errors, and 17 (9 high, 8 low) for omission errors.

#### 2.5.3 LZSUM and Regions of Interest

To evaluate the complexity level of each participant for both levels of arousal, LZSUM values were averaged across trials and electrodes for both alertness and drowsiness. Next, we performed two-way mixed factor ANOVAs to analyse the interaction between level of arousal (alert/drowsy) as the within-subject factor, performance (high/low) as the between-subject factor, and LZSUM as the dependent variable. To further investigate the potential interaction between level of arousal and performance, we conducted the above-described analysis again, but now with one extra factor: regions of interest (ROIs). Using the same approach as described in Folland et al. (2015), we divided 90 of the 129 electrodes into five areas (frontal, central, temporal, parietal, and occipital) for both the left and right hemispheres, which resulted in 10 ROIs in total. Each ROI consisted of approximately 16 to 20 electrodes, then averaged to represent the EEG activity from that brain area. Midline electrodes were excluded to aid an accurate comparison of the left and right hemispheres. Also, the outermost electrodes were excluded to avoid interference from facial muscle artefacts. Next, LZSUM was computed for each ROI, and per condition (alert/drowsy). Finally, we performed three-way mixed factor ANOVAs to analyse the interaction between level of arousal (alert/drowsy) and ROI as the within-subject factors, performance (high/low) as the between-subject factor, and LZSUM as the dependent variable.

## 3. RESULTS

### 3.1 GROUP COMPARISONS

First, we examined performance from a group-level perspective. Participants were categorised into high and low performers to investigate the relationship between group-level performance, LZSUM (both whole brain and ROI-based), and levels of arousal (drowsy vs. alert).

#### 3.1.1 LZSUM in High and Low Performers Across Arousal States

We performed a two-way mixed factors ANOVA for each of the 4 performance metrics (mean RT, omission errors, commission errors, and CV RT) to examine the interaction between arousal level (drowsy, alert) and performance (high, low) with LZSUM as the outcome variable. This analysis aimed to investigate differences in informational complexity between high and low performers during different states of arousal. In this way, we aimed to study LZSUM as a potential neural marker for the underlying compensatory mechanism that allows high performers to maintain their performance level while under neural strain. Our results show a significant interaction effect (performance x arousal) for mean RT [*F*(1,21) = 4.82, *p* = .040, *ηp*^*2*^ = 0.19], depicted in figure 2a, and omission errors [*F*(1,15) = 5.81, *p* = .029, *ηp*^*2*^= 0.28], shown in figure 2b. However, no significant interaction effect was found for commission errors [*F*(1,17) = 1.34, *p* = .263, *ηp*^*2*^ = 0.07] nor CV RT [*F*(1,21) = 0.43, *p* = .518, *ηp*^*2*^ = 0.02]. Further post-hoc t-tests showed that high performers exhibited a significant increase in LZSUM when drowsy compared to when alert. This was the case for both mean RT [*t*(10) = -3.54, *p* = .005, Cohen’s *d* = - 1.12] and omission errors [*t*(8) = -2.98, *p* = .018, Cohen’s *d* = -1.06]. Conversely, no significant differences were found between drowsy and alert states for low performers in terms of mean RT [*t*(11) = -0.16, *p* = .870, Cohen’s *d* = -0.05] and omission errors [*t*(7) = 0.45, *p* = .660, Cohen’s *d* = 0.17]. We also conducted t-tests to assess the LZSUM differences between high and low performers in each arousal state. These showed that compared to low performers, high performers exhibit significantly higher LZSUM values in the alert state for both mean RT [*t*(21) = 2,92, *p* = .008, Cohen’s *d* = 1.28] and omission errors [*t*(15) = 3.47, *p* = 0.003, Cohen’s *d* = 1.79], and in the drowsy state for both mean RT [*t*(21) = 3.95, *p* < .001, Cohen’s *d* = 1.72] and omission errors [*t*(15) = 5.13, *p* < .001, Cohen’s *d* = 2.65]. These results show that the differences between the high performing and low performing group became more apparent in the drowsy state. Furthermore, LZSUM appeared to be higher overall in high performers compared to low performers for both mean RT [*t*(44) = 4.91, *p* < .001, Cohen’s *d* = 1.48] and omission errors [*t*(32) = 6.09, *p* < .001, Cohen’s *d* = 2.16]. The observed significant increase in LZSUM for high performers while drowsy contrasted with the absence of this effect in low performers, suggesting an underlying compensatory mechanism that enabled high performers to maintain their performance levels despite the neural strain induced by drowsiness.

**Figure 1.**
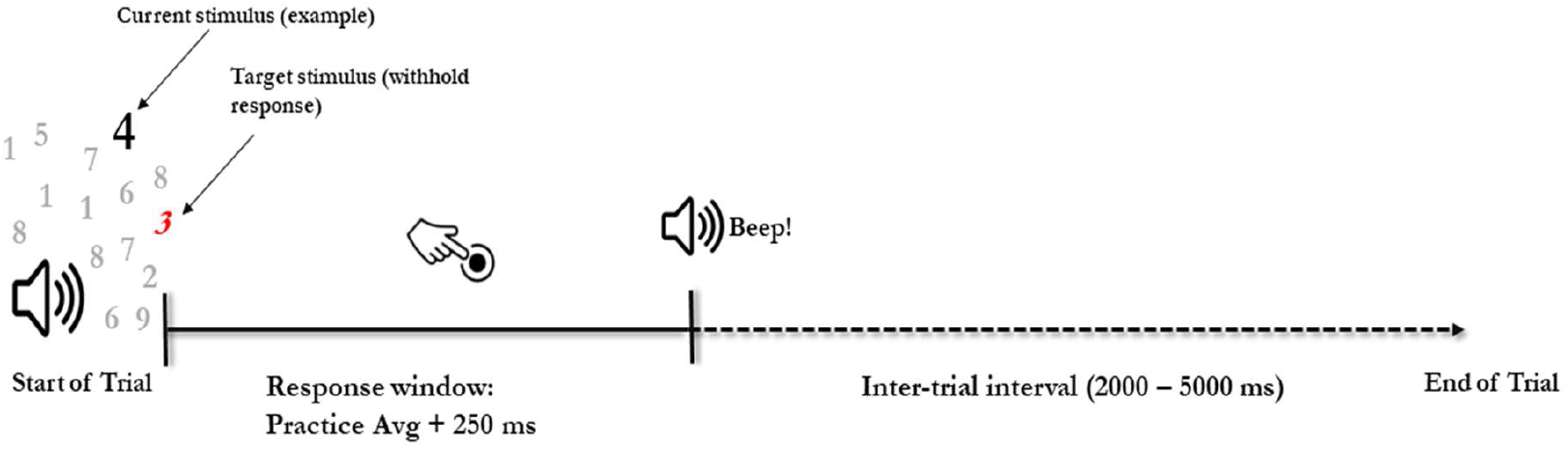
Single-trial structure of the auditory version of the Sustained Attention to Response Task (SART). At the start of each trial, a recording was played of a number between 0 and 9 (e.g. 4). The participant had to press a button after hearing the number, except when hearing the number 3 (i.e. target stimulus) when a response should be withheld. The length of the response window was determined for each participant by their average response time during the practice run plus 250ms. The end of the response window was indicated with a beep followed by a random inter-trial interval of 2 to 5 seconds.

**Figure 2.**
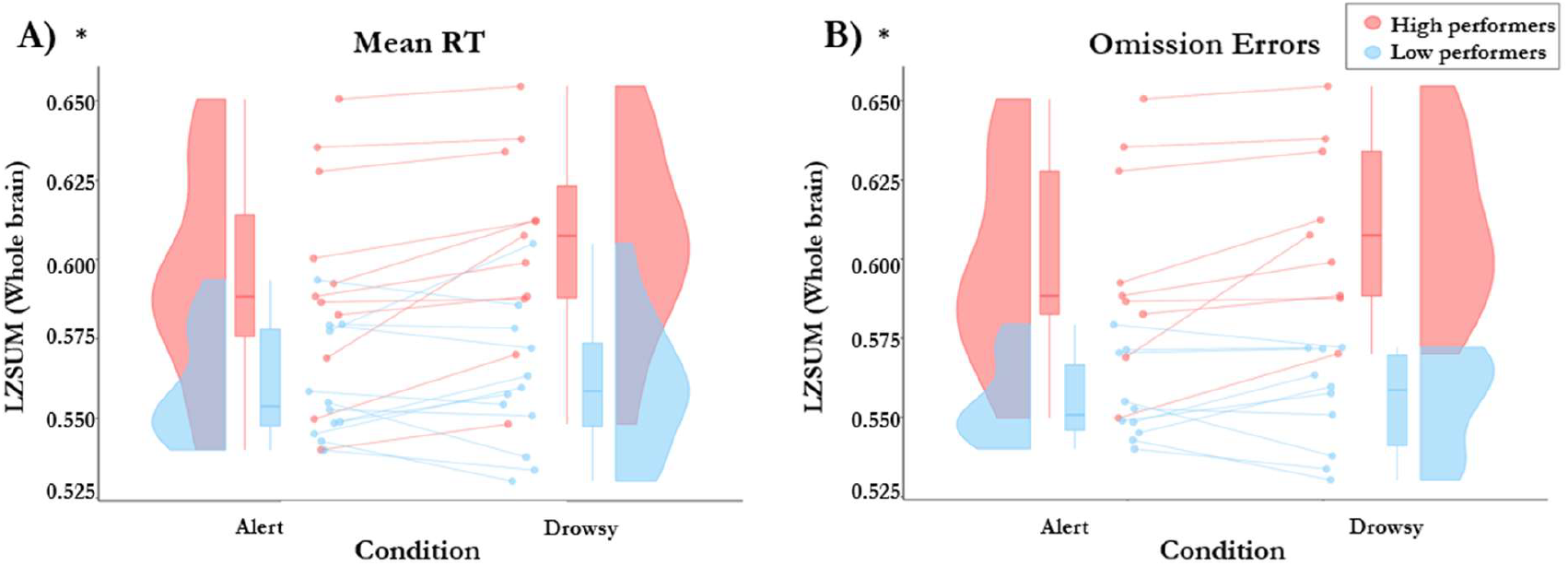
LZSUM in alert and drowsy states: comparing high to low performers. **2A**. This panel displays a rain cloud plot showing the variations of LZSUM values derived from whole-brain EEG signals across alert and drowsy conditions. The x-axis represents the two conditions, and the y-axis represents the LZSUM values. Participants are divided into two groups based on their mean reaction times (RT) when performing the auditory version of the Sustained Attention to Response Task (SART). High performers are represented in red and low performers in blue. **2B**. Similar to panel 2A, but here, participants are categorised into high and low performers based on the number of omission errors when performing the SART. (**p ≤ 0.05 *), (p ≤ 0.01**), (p ≤ 0.001***)**

#### 3.1.2 Region of Interest Analysis of LZSUM in High and Low Performers Across Arousal States

We conducted three-way ANOVA analyses to examine the interaction between performance, level of arousal, and region of interest (ROI) with LZSUM as the outcome variable. In this way, we aimed to investigate which ROIs are most involved in the effect that showed LZSUM differences when comparing high performers and low performers across different levels of arousal. Notable variations were observed across different performance metrics. For CV RT, there was a highly significant three-way interaction effect [*F*(9,189) = 5.89, *p* < .001, ηp^2^ = 0.22], as well as a significant two-way interaction between performance and ROI [*F*(9,189) = 5.09, *p* < .001, ηp^2^ = 0.20]. However, no significant interaction was found between performance and arousal [*F*(1,21) = 0.32, *p* = .578, *ηp*^*2*^= 0.02], and the main effect of performance was not significant [*F*(1,21) = 2.76, *p* = .112, *ηp*^*2*^= 0.12]. In the case of mean RT, a moderate but significant three-way interaction was noted [*F*(9, 89) = 2, *p* = .041, *ηp*^*2*^= 0.09], alongside a significant two-way interaction between performance and ROI [*F*(9, 89) = 2.09, *p* = .033, *ηp*^*2*^= 0.09]. The interaction between performance and arousal showed a trend with *F* (1,21) = 3.5, *p* = .075, ηp^2^= 0.14], and the main effect of performance was significant [*F*(1,21) = 7.28, *p* = .013, *ηp*^*2*^= 0.26]. For omission errors, the three-way interaction was not significant [*F*(9, 135) = 0.81, *p* = .609, *ηp*^*2*^= 0.05]. However, a significant two-way interaction was observed for performance * ROI [*F*(9, 135) = 2.89, *p* = .001, *ηp*^*2*^= 0.16], and a trend for performance * arousal [*F*(1, 15) = 4.35, *p* = .055, ηp^2^= 0.22), and the main effect of performance was significant [*F*(1, 15) = 11.55, *p* = 0.004, *ηp*^*2*^= 0.44]. For commission errors, the three-way interaction was significant [*F*(9, 153) = 2.01, *p* = .041, *ηp*^*2*^= 0.11], whereas no significant two-way interactions or main effects were observed for performance * ROI [*F*(9, 153) = 0.96, *p* = .478, *ηp*^*2*^= 0.05], performance * arousal [*F*(1, 17) = 0.81, *p* = .380, *ηp*^*2*^= 0.05], or the main effect of performance [*F*(1, 17) = 0.93, *p* = .347, *ηp*^*2*^= 0.05].

Subsequent post-hoc analyses examining the interaction between arousal condition and performance across specific ROIs revealed significant effects after Bonferroni correction. In the frontal left (FL) region, the condition by performance interaction for mean RT was significant [*F* (1, 21) = 12, *p* = .023, *ηp*^*2*^ = 0.36], as was the interaction in the frontal right (FR) region [*F*(1, 21) = 11.93, *p* = .024, *ηp*^*2*^ = 0.36]. Similarly, for omission errors, significant interactions were observed in both the FL [*F*(1, 15) = 19.2, *p* = 0.005, *ηp*^*2*^ = 0.56] and FR [*F*(1, 15) = 15.2, *p* = .014, *ηp*^*2*^ = 0.50], as well as in the central left (CL) region [*F*(1, 15) = 15.22, *p* = .014, *ηp*^*2*^ = 0.50]. Noteworthy trends after correction were also found for mean RT in both the CL [*F*(1, 21) = 8.33, *p* = .089, *ηp*^*2*^=0.28] and CR [*F*(1, 21) = 8.06, *p* = .098, *ηp*^*2*^=0.28] and for omission errors in the CR [*F*(1, 15) = 10.06, *p* = .063, *ηp*^*2*^= 0.40]. No significant results or trends were observed for CV RT or commission errors, or in any of the other ROIs. See figure 3 for an overview of these region-specific effects.

**Figure 3.**
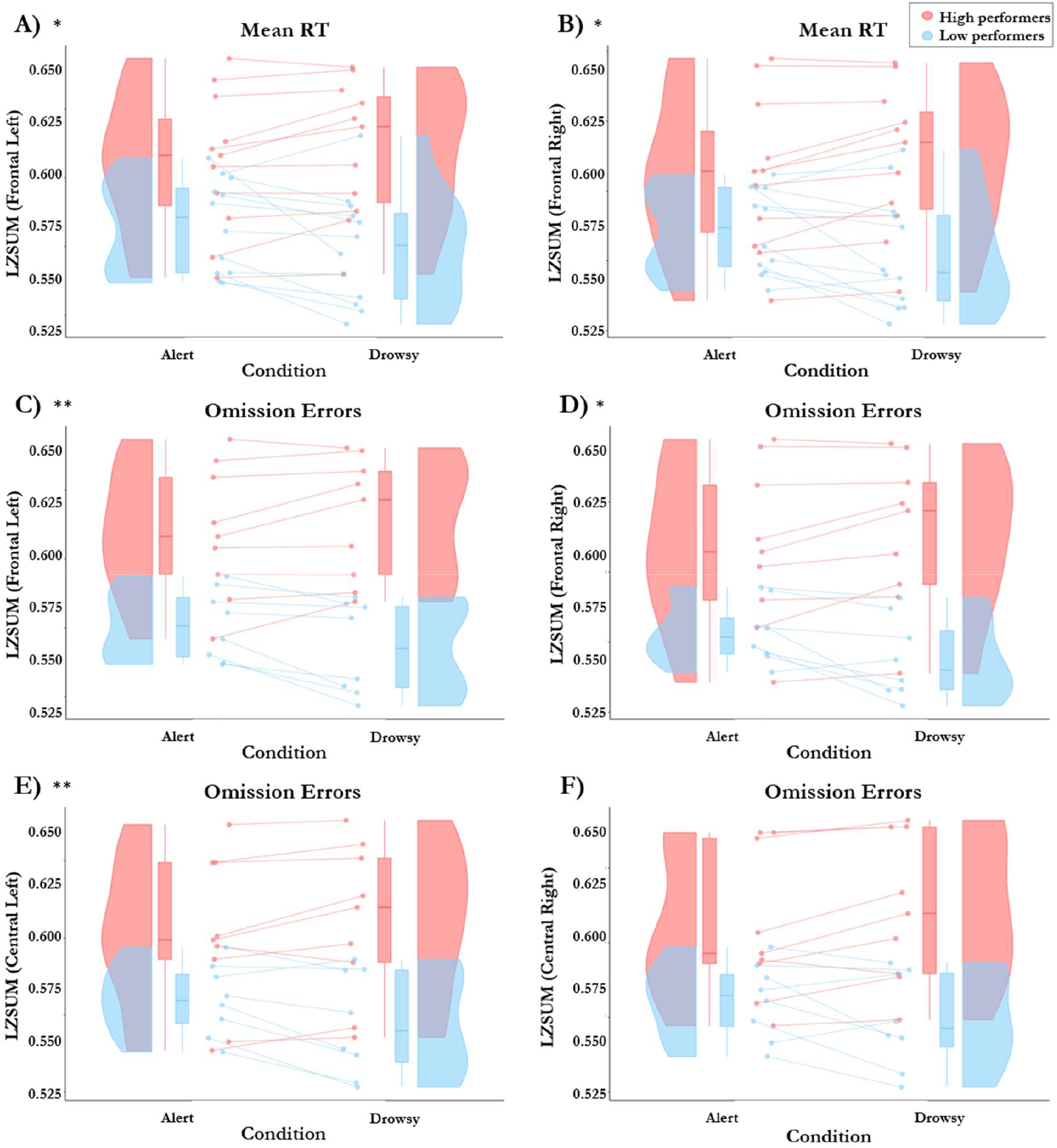
LZSUM ROI-based EEG analysis (LZSUM) in alert and drowsy states: comparing high to low performers. All panels display rain cloud plots showing the variations of LZSUM values derived from various regions of interest (ROIs) (frontal, central, temporal, parietal, and occipital) across alert and drowsy conditions. Only the significant results after Bonferroni correction are included in the panels. The x-axis represents the two conditions (alert vs. drowsy), and the y-axis represents the LZSUM values. Participants are divided into two groups based on their performance on the auditory version of the Sustained Attention to Response Task (SART). High performers are represented in red and low performers in blue. **3A**. Variations of LZSUM values derived from *frontal left* EEG signals across alert and drowsy conditions. Performance is determined by mean reaction times (RT) **3B**. Similar to 3A, but here, LZSUM values were derived from *frontal right* EEG signals. **3C**. Variations of LZSUM values derived from *frontal left* EEG signals across alert and drowsy conditions. Performance is determined by omission errors. **3D**. Similar to 3C, but here, LZSUM values were derived from *frontal right* EEG signals. **3E**. Variations of LZSUM values derived from *central left* EEG signals across alert and drowsy conditions. **3F**. Similar to 3E, but here, LZSUM values were derived from *central right* EEG signals. (**p ≤ 0.05 *), (p ≤ 0.01**), (p ≤ 0.001***). Note:** Significance asterisks reflect **corrected** p-values.

Further analysis using post-hoc t-tests on these significant interactions indicated that in the FL region, high performers showed a significant increase in LZSUM values when drowsy for mean reaction time [*t*(10) = -2.76, *p* = .020, Cohen’s *d* = -0.87] and omission errors [*t*(8) = -2.34, *p* = .047, Cohen’s *d* = - 0.83], whereas low performers exhibited significant decreases [mean RT: *t*(11) = 2.54, *p* = .027, Cohen’s *d* = 0.77; omission errors: *t*(7) = 3.84, *p* = .006, Cohen’s *d* = 1.45]. A similar pattern was observed in the FR region, with high performers experiencing a significant increase in LZSUM values when drowsy [mean RT: *t*(10) = -3.13, *p* = .010, Cohen’s *d* = -0.99; omission errors: *t*(8) = -2.51, *p* = .037, Cohen’s *d* = -0.89] and low performers showing a significant decrease [mean RT: *t*(11) = 2.34, *p* = .039, Cohen’s *d* = 0.71; omission errors: *t*(7) = 2.91, *p* = .023, Cohen’s *d* = 1.10]. In the CL, high performers also had a significant increase in LZSUM values for omission errors when drowsy [*t*(8) = -2.46, *p* = 0.040, Cohen’s *d* = -0.87], with low performers again showing a significant decrease [*t*(7) = 2.96, *p* = .021, Cohen’s *d* = 1.12]. These results suggest a region-dependent difference in the modulating effect of arousal states on LZSUM when comparing high performers to low performers, particularly concerning the performance in terms of mean RT and omission errors. The effect is most pronounced in key regions associated with attentional and executive processes (frontal regions) (Kievit et al., 2014) and areas involved in sensorimotor functions (central regions) (Kandel et al., 2013). This pattern suggests a similar compensatory mechanism as was also observed in the whole-brain LZSUM results supported by a region-specific modulation of brain complexity. It must be mentioned that since EEG has limited spatial precision (Nunez & Srinivasan, 2006), effects found in specific regions may originate from neighbouring areas. However, the significant effect found in the frontal and central regions, and the absence of an effect in other regions (e.g. the occipital electrodes), suggests meaningful region-dependent activity.

### 3.2 CORRELATIONAL ANALYSIS OF TASK PERFORMANCE WITH BRAIN MEASURES AND COGNITIVE RESERVE PROXIES

We further explored the relationship between task performance and key variables such as white matter network connectivity and proxies of cognitive reserve with a correlational analysis. We calculated correlations between the four metrics of task performance (mean RT, CV RT, commission errors, and omission errors) and: 1) LZSUM whole brain and in each of the 10 ROIs, 2) graph theoretical measures of white matter connectivity, 3) grey matter volume in cortical and hippocampal areas, and 4) proxies of cognitive reserve

#### 3.2.1 Correlational analysis of LZSUM and task performance

Building on our earlier findings that showed a strong effect for performance on LZSUM differentiating between high and low performers, we now perform the corresponding individual differences analysis. Significant negative correlations were found between various performance metrics and LZSUM in the frontal and central areas, the temporal right area, and LZSUM whole brain. The most robust negative correlations, which remained statistically significant after Bonferroni correction, involved LZSUM in frontal and central regions. These results support our previous findings by demonstrating a consistent pattern where LZSUM negatively correlates with task performance difference (alert vs. drowsy), especially in frontal and central brain areas. Here, a higher LZSUM ratio (drowsy/alert) is associated with a better ability to maintain task performance under drowsiness, as indicated by lower scores in performance difference (drowsy/alert for RT, drowsy – alert for errors), and vice versa. This negative correlation is mainly prominent in frontal and central brain areas, indicating that increased LZSUM in these areas during drowsiness is linked to more resilient cognitive functioning under neural strain. For a detailed overview of all significant correlations, please refer to figure 4 below or Table S3 in the supplementary materials.

**Figure 4.**
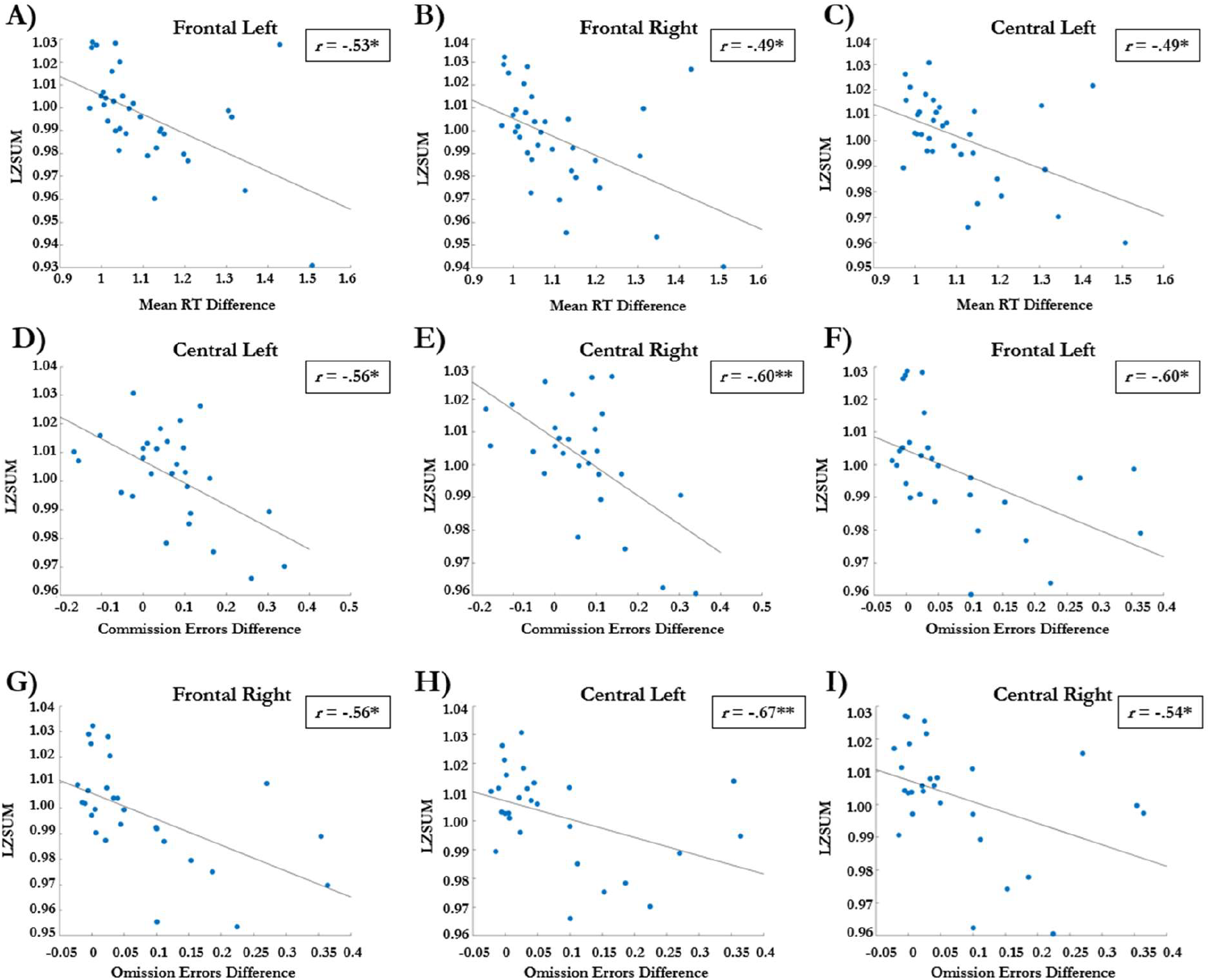
Scatterplots of associations between LZSUM and performance difference (drowsy vs alert) on various performance metrics. All panels display scatterplots which illustrate the significant relationship between LZSUM of specific regions of interest (ROI) (y-axis) and various cognitive performance metrics (x-axis). All of the significant results displayed in the scatterplots survived Bonferroni correction. **4A**. Association between frontal left LZSUM and mean RT difference. **4B**. Association between frontal right LZSUM and mean RT difference. **4C**. Association between central left LZSUM and mean RT difference. **4D** Association between central left LZSUM and commission errors difference. **6E**. Association between central right LZSUM and commission errors difference. **4F**. Association between frontal left LZSUM and omission errors difference. **5G**. Association between frontal right LZSUM and omission errors difference. **4H**. Association between central left LZSUM and omission errors difference. **4I**. Association between central right LZSUM and omission errors difference. (**p ≤ 0.05 *), (p ≤ 0.01**), (p ≤ 0.001***). Note:** Significance asterisks reflect **corrected** p-values. Only the correlations that survived correction were plotted here.

#### 3.2.2 Correlational analysis of structural brain measures, cognitive reserve proxies, and task performance

We computed network measures of white matter connectivity, using the Brain Connectivity Toolbox (Rubinov & Sporns, 2010), to investigate various network characteristics, including characteristic path length, average degree, global efficiency, mean clustering coefficient, mean participation coefficient, small-worldness, network density, and core-periphery structure. For a detailed overview of these measures in the context of cognition, see Rubinov and Sporns (2010). These metrics were calculated for both the overall brain network and seven distinct subnetworks as defined by Yeo et al. (2011), using the Brain Connectivity Toolbox (Rubinov & Sporns, 2010). Additionally, our analysis included structural volumetric MRI data of cortical and medial temporal lobe (MTL) regions. To further contextualise our findings within the framework of cognitive reserve, we included various proxies of cognitive reserve, namely two distinct IQ measures: Cattell (Cattell, 1973) and NART (Nelson & Willison, 1991) along with the discrepancy between them (NART – Cattell), the scores on the Montreal Cognitive Assessment (MoCA) (Nasreddine, 2005), and educational level.

We calculated correlations between the four metrics of behavioural performance (mean RT, CV RT, commission errors, and omission errors) and: 1) network measures of white matter connectivity, 2) grey matter volume in cortical areas and the MTL, and 3) proxies of cognitive reserve. Our results did not reveal any significant correlations for the proxies of cognitive reserve, however, various significant correlations were observed between behavioural performance, white matter connectivity, and grey matter volume. However, these did not survive Bonferroni correction. These findings are discussed in more detail in the Supplementary Materials (Sections S1-S3, Tables S1-S2, and Figures S1-S2).

## 4. DISCUSSION

The primary goal of the current work was to gain more insight into the underlying neural mechanisms of cognitive reserve in an elderly population. We provided an operational definition and simple method aimed at measuring cognitive reserve more directly and related this to a variety of potential structural and functional neural markers. Here, cognitive reserve was operationalised as the performance discrepancy (i.e. reaction time, errors) between drowsy and alert states on a sustained attention task. We hypothesised that the informational complexity of the EEG signal, as quantified by Lempel-Ziv complexity for single channels (LZSUM), would decrease in low performers when transitioning from an alert to a drowsy state, but would be maintained or increased in high performers, reflecting a compensatory mechanism.

The main findings from our study show that in high performing individuals LZSUM increases when transitioning from an alert to a drowsy state, which contrasted with the pattern observed in low performers, who showed no significant change in LZSUM under similar conditions. These LZSUM effects were primarily observed in the frontal and central regions demonstrating that there are region-dependent differences in the modulating effect of arousal on LZSUM, with the effect most pronounced in regions related to attention, executive (frontal) (Kievit et al., 2014), and sensorimotor (central) functioning (Kandel et al., 2013). This finding is consistent with Boncompte et al. (2021), who reported an increase in Lempel-Ziv complexity during propofol-induced mild sedation in participants maintaining behavioural performance, which was observed in the frontal electrodes. Taken together, increases in informational complexity under neural strain in the frontal areas may predict higher levels of cognitive reserve, supported by the potential involvement of the fronto-parietal control network in cognitive reserve (Kievit et al., 2014; Stern et al., 2019). However, these findings need to be interpreted with caution since the limited spatial precision of EEG means that effects found in specific regions may originate from neighbouring areas (Nunez & Srinivasan, 2006). However, the significant effect found in the frontal and central regions, and the absence of an effect in other regions (e.g. the occipital electrodes), suggests meaningful region-dependent activity. Future research using neuroimaging methods with higher spatial precision such as fMRI could complement our results and would aid in a more detailed localisation of the effect.

Additionally, in concert with the above results, the LZSUM correlation results showed that a higher LZSUM ratio (drowsy/alert) is associated with smaller performance differences (drowsy/alert for reaction time, drowsy – alert for errors), indicating a better ability to maintain task performance under drowsiness, with high performers showing minimal decreases or slight improvements, and low performers experiencing a more substantial decline in performance.

Finally, our results showed LZSUM values to be higher in high performers overall, independent of the level of arousal, and this effect could be more clearly observed during drowsiness. The consistent pattern across different states of arousal and brain regions that differentiates high performers from low performers highlights the potential role of individual differences in informational complexity as a neural marker of cognitive reserve.

We also measured correlations between our measure of cognitive reserve (i.e. performance difference alert vs. drowsy) and well-known proxies of cognitive reserve, such as IQ and educational level. No significant correlations were found which can be explained in various ways. It could be that the statistical power of the study was too low to uncover the expected association between the proxies of cognitive reserve and ability to maintain task performance under neural strain. LZSUM may be a more sensitive measure so that even with low statistical power a strong effect could still be found, but the relatively small sample size may have limited the detection of more subtle effects. Future research with larger samples could offer richer distributions and further validate our findings. Additionally, other well-known proxies were not included in the study such as current lifestyle and activity levels, social life and leisure activities, and occupational complexity (Satz, 1993; Scarmeas et al., 2001; Valenzuela et al., 2008). Proxies such as premorbid IQ and educational level may not fully capture the current contribution to cognitive reserve. Factors such as current leisure activities, social life and activity level may be of more influence in the elderly, as these relate to current ongoing physical and mental engagement and could be included in future studies.

At this stage, we cannot make definitive conclusions about whether we have measured cognitive reserve. Our main result that shows informational complexity differences between high and low performers could potentially be attributed to task-specific effects, such as the ability to compensate for neural strain on a sustained attention task. However, a similar effect of informational complexity differences between high and low performers was found in the Boncompte et al. (2021) study, where they used an auditory discrimination task. Auditory sustained attention tasks and auditory discrimination tasks differ in specific demands (sustained attention versus perceptual discrimination), suggesting the effect may be reflecting a broader cognitive effect, instead of an effect that is purely task-specific. To further investigate whether our finding relates to a broader cognitive compensatory mechanism, the experiment could be replicated using a variety of different tasks related to different perceptual modalities, different types of cognitive functioning (e.g. memory) and tasks that are more cognitively demanding (e.g. n-back task).

Another important potential limitation to consider is that perhaps our group of participants were too homogeneous in terms of proxies cognitive reserve. For instance, on average our participants had spent 18 years in education. In previous research that assessed the relationship between educational attainment and cognitive reserve, low educational attainment is defined as spending 8 years or less in education (Valenzuela & Sachdev, 2006). Since our group consisted primarily of highly educated participants it may be that a threshold was reached beyond which additional years of education yield diminishing returns in terms of further increasing cognitive reserve.

A study with the same drowsiness paradigm in patients with mild cognitive impairment (MCI) and Alzheimer’s disease (AD) may lead to additional insights of whether we are truly measuring cognitive reserve. High cognitive reserve individuals with AD or MCI are characterised by relatively well-preserved cognitive functioning given their advanced brain pathology (Steffener & Stern, 2012). Therefore, if we compare healthy elderly participants with patients with MCI and AD, using objective measures of brain pathology (e.g. structural MRI) alongside cognitive functioning assessments, we can more accurately differentiate between high and low cognitive reserve patients. This clearer differentiation could further our understanding of the experimental effects and potentially validate the findings of the current work.

We also explored correlations between performance difference (alert vs. drowsy) and various measures, including cortical and hippocampal grey matter volume and white matter connectivity. Volumetric changes in various key brain regions were associated with all performance metrics besides omission errors. For instance, there was a positive association between volume in the left entorhinal cortex and making more commission errors while drowsy. Of chief importance are frontal, parietal and medial temporal lobe (MTL) related brain regions because of their involvement in Alzheimer’s Disease (AD). These regions are crucial as they are implicated in core functions such as memory, spatial orientation, and executive function, which deteriorate in AD (Jack et al., 2010).

It is worth noting that associations between volumetric variations in the medial temporal lobe regions and the ability to maintain performance under neural strain are of potential clinical significance as these brain regions are among the earliest to atrophy in AD (Jack et al., 2010). The observation that lower volume in the entorhinal cortex correlates with increased ability to maintain performance while drowsy potentially underscores the concept of cognitive reserve. This counterintuitive finding suggests that individuals with higher cognitive reserve may effectively utilise alternate neural pathways to maintain cognitive performance despite structural brain changes, indicating a complex interaction between the beginning stages of neural degeneration and compensatory mechanisms. However, none of the volumetric findings survived correction for multiple comparisons.

Furthermore, our network analysis of white matter connectivity showed some interesting associations, such as a positive correlation between small-worldness in the frontal-parietal control network and omission errors, but similarly to the volumetric analysis none of the correlations survived correction for multiple comparisons. Importantly, we used an exploratory approach by including a high number of variables to uncover potentially meaningful associations. However, a relatively small sample size combined with many comparisons may have reduced statistical power and potentially caused some genuine effects to be missed. Therefore, any significant correlations before correction should be seen as preliminary and warrant further investigation into the role of grey matter volume, as well as network characteristics, in cognitive reserve. However, the combination of the inconclusiveness of the structural brain metrics and proxies of cognitive reserve in combination with the strong results we found for LZSUM suggests that complexity measures such as LZSUM derived from electrophysiological data during a task are statistically robust and capture unique information that various other metrics do not. Thus, we argue that LZSUM is a strong neural candidate to index cognitive reserve.

Interestingly, measures of informational complexity such as Lempel-Ziv are used extensively in consciousness research. Informational complexity measures have been shown to accurately assess anaesthetic depth (Schartner et al., 2015; Shin et al., 2020), differentiate between levels of consciousness in brain-injured patients (Chennu et al., 2014; King et al., 2013), and are useful to study transitions in sleep stages (Casali et al., 2013), and show increased values in studies with psychedelics (Schartner et al., 2017). These studies all show a similar pattern: lower levels of consciousness correspond to lower informational complexity, and vice versa. Critically, our findings present a counterexample to the prevailing view that informational complexity purely reflects conscious level. Given that in the high-performing group, informational complexity dissociated from conscious state, our study challenges how complexity measures are interpreted in consciousness research. Our results suggest that the increase in complexity might be related to the brain’s capacity to adapt and compensate under varying cognitive demands, similar to other studies such as the work from Boncompte et al. (2021) that explore subtler fluctuations in consciousness (e.g. moderate sedation, drowsiness), where informational complexity shows a more non-linear pattern. In this way, our work invites a re-evaluation of how informational complexity is interpreted in consciousness research.

In conclusion, the examination of differences in performance from alert to drowsy states as an operational measure of cognitive reserve offers a novel approach compared to traditional proxies such as educational level and IQ. This method directly captures the active process of compensating for neural challenges, which is more representative of the real-world scenarios faced by individuals with cognitive decline, and in this way potential underlying neural markers of cognitive reserve can be investigated. Our findings show potential for LZSUM as a neural marker in cognitive reserve assessment. Future validation of these findings could lead to its use in clinical trials, particularly in evaluating cognitive reserve as a factor in treatment response. For instance, assessing cognitive reserve using LZSUM in clinical trials for Alzheimer’s disease medications could help control for cognitive reserve as a confounding factor, leading to more precise and personalised treatment strategies. Such applications could significantly advance the field of neurodegenerative disease treatment, offering new tools for clinicians in their efforts to mitigate the impact of cognitive decline.

## Supplementary data

Further analyses exploring the associations between task performance under different states of alertness and various brain structural characteristics, as well as known proxies of cognitive reserve, were conducted. These results are discussed in more detail in the Supplementary Materials (Sections S1-S3, Tables S1-S2, and Figures S1-S2).

## Data availability

The main and supplementary data including data dictionary can be found on https://osf.io/w8qtn/

## CRediT authorship contribution statement

**Laura Stolp:** Conceptualisation, Investigation, Formal Analysis, Data Curation, Visualisation, Writing – Original Draft, Writing – Review & Editing, **Kanad N Mandke:** Conceptualisation, Investigation, Supervision, Writing – Review & Editing, **Pedro AM Mediano:** DTI Preprocessing, Resources, **Helena M Gellersen:** Data Collection, MRI Preprocessing, Writing – Review & Editing, **Alex Swartz:** Investigation, Data Collection, **Katarzyna Rudzka:** Investigation, Data Collection, **Jon Simons:** Resources, **Tristan A Bekinschtein:** Conceptualisation, Investigation, Resources, Writing – Review & Editing, **Daniel Bor:** Conceptualisation, Investigation, Writing – Review & Editing, Supervision, Project Administration, Funding Acquisition

## Acknowledgements

We would like thank Sridhar R. Jagannathan for his assistance with the drowsiness classification of the EEG signal. We also extend our appreciation to all the participants who volunteered to take part in the study, as well as to all departments involved across the University of Cambridge for facilitating the study’s set-up and delivery.

## REFERENCES

Bassett, D. S., & Sporns, O. (2017). Network neuroscience. Nature neuroscience, 20(3), 353–364. 10.1038/nn.4502

Braak, H., & Braak, E. (1991). Neuropathological stageing of Alzheimer-related changes. Acta neuropathologica, 82(4), 239–259. 10.1007/bf00308809

Bigdely-Shamlo, N., Mullen, T., Kothe, C., Su, K. M., & Robbins, K. A. (2015). The PREP pipeline: standardized preprocessing for large-scale EEG analysis. Frontiers in neuroinformatics, 9, 16. 10.3389/fninf.2015.00016

Blondel, V. D., Guillaume, J. L., Lambiotte, R., & Lefebvre, E. (2008). Fast unfolding of communities in large networks. Journal of statistical mechanics: theory and experiment, 2008(10). 10.1088/1742-5468/2008/10/P10008

Boncompte, G., Medel, V., Cortínez, L. I., & Ossandón, T. (2021). Brain activity complexity has a nonlinear relation to the level of propofol sedation. British Journal of Anaesthesia, 127(2), 254–263. 10.1016/j.bja.2021.04.023

Carr, V. A., Bernstein, J. D., Favila, S. E., Rutt, B. K., Kerchner, G. A., & Wagner, A. D. (2017). Individual differences in associative memory among older adults explained by hippocampal subfield structure and function. Proceedings of the National Academy of Sciences, 114(45), 12075–12080. 10.1073/pnas.171330811

Casali, A.G., Gosseries, O., Rosanova, M., Boly, M., Sarasso, S., Casali, K.R., Casarotto, S., Bruno, M., Laureys, S., Tononi, G., & Massimini, M. (2013). A theoretically based index of consciousness independent of sensory processing and behavior. Science Translational Medicine, 5(198), 10.1126/scitranslmed.3006294

Cattell, R. B. (1973). Culture-fair intelligence test. Institute for Personality and Ability Testing. 10.1037/t14354-000

Chennu, S., Finoia, P., Kamau, E., Allanson, J., Williams, G. B., Monti, M. M., … & Cabezas-Soto, D. (2014). Spectral signatures of reorganised brain networks in disorders of consciousness. PLoS computational biology, 10(10). 10.1371/journal.pcbi.1003887

Cummings, J., Ritter, A., & Zhong, K. (2018). Clinical trials for disease-modifying therapies in Alzheimer’s disease: a primer, lessons learned, and a blueprint for the future. Journal of Alzheimer’s Disease, 64(1), S3–S22. doi: 10.3233/JAD-179901

Delorme, A., & Makeig, S. (2004). EEGLAB: an open source toolbox for analysis of single-trial EEG dynamics including independent component analysis. Journal of neuroscience methods, 134(1), 9–21. 10.1016/j.jneumeth.2003.10.009

Delorme, A., Majumdar, A., Sivagnanam, S., Martinez-Cancino, R., Yoshimoto, K., & Makeig, S. (2019, March). The open EEGLAB portal. In 2019 9th International IEEE/EMBS Conference on Neural Engineering (NER) (pp. 1142–1145). IEEE. 10.1109/ner.2019.8717114

Desikan, R. S., Ségonne, F., Fischl, B., Quinn, B. T., Dickerson, B. C., Blacker, D., … & Killiany, R. J. (2006). An automated labeling system for subdividing the human cerebral cortex on MRI scans into gyral based regions of interest. Neuroimage, 31(3), 968–980. 10.1016/j.neuroimage.2006.01.021

Durmer, J. S., & Dinges, D. F. (2005, March). Neurocognitive consequences of sleep deprivation. In Seminars in neurology (Vol. 25, No. 01, pp. 117–129. 10.1055/s-2005-867080

Duthey, B. (2013). Priority Medicines for Europe and the World. “A public health approach to innovation”. WHO Background paper, 6.11, Alzheimer Disease and other Dementias.

Fischl, B., & Dale, A. M. (2000). Measuring the thickness of the human cerebral cortex from magnetic resonance images. Proceedings of the National Academy of Sciences, 97(20), 11050–11055. 10.1073/pnas.200033797

Folland, N. A. etal. (2015). Journal of cognitive neuroscience, 27(5), 1060–1067. 10.1162/jocn_a_00764

Fratiglioni, L., & Wang, H. X. (2007). Brain reserve hypothesis in dementia. Journal of Alzheimer’s disease, 12(1), 11–22. 10.3233/JAD-2007-12103

Gautam, P., Cherbuin, N., Sachdev, P. S., Wen, W., & Anstey, K. J. (2011). Relationships between cognitive function and frontal grey matter volumes and thickness in middle aged and early old-aged adults: The PATH Through Life Study. Neuroimage, 55(3), 845–855. 10.1016/j.neuroimage.2011.01.015

Gellersen, H. M., Trelle, A. N., Farrar, B. G., Coughlan, G., Korkki, S. M., Henson, R. N., & Simons, J. S. (2023). Medial temporal lobe structure, mnemonic and perceptual discrimination in healthy older adults and those at risk for mild cognitive impairment. Neurobiology of Aging, 122, 88–106. 10.1016/j.neurobiolaging.2022.11.004

Goupil, L., & Bekinschtein, T. (2012). Cognitive processing during the transition to sleep. Archives italiennes de biologie, 150(2/3), 140–154. 10.4449/aib.v150i2.1247

Huang, L. K., Kuan, Y. C., Lin, H. W., & Hu, C. J. (2023). Clinical trials of new drugs for Alzheimer disease: a 2020–2023 update. Journal of Biomedical Science, 30(1), 83. 10.1186/s12929-023-00976-6

Iber, C., Ancoli-Israel, S., Chesson, A. L., & Quan, S. F. (2007). The AASM manual for the scoring of sleep and associated events: rules, terminology and technical specifications (Vol. 1). Westchester, IL: American academy of sleep medicine. 10.5664/jcsm.26812

Jack, C. R., Knopman, D. S., Jagust, W. J., Shaw, L. M., Aisen, P. S., Weiner, M. W., … & Trojanowski, J. Q. (2010). Hypothetical model of dynamic biomarkers of the Alzheimer’s pathological cascade. The Lancet Neurology, 9(1), 119–128. 10.1016/s1474-4422(09)70299-6

Jagannathan, S. R., Nassar, A.E., Jachs, B., Pustovaya, O.V., Bareham, C.A., & Bekinschtein, T.A. (2018). Tracking wakefulness as it fades: micro-measures of Alertness. NeuroImage, 176, 138–151. 10.1016/j.neuroimage.2018.04.046

Jones, R. N., Manly, J., Glymour, M. M., Rentz, D. M., Jefferson, A. L., & Stern, Y. (2011). Conceptual and measurement challenges in research on cognitive reserve. Journal of the International Neuropsychological Society, 17(4), 593–601. 10.1017/s1355617710001748

Ju, Y. E., Lucey, B.P., & Holtzman, D.M. (2014). Sleep and Alzheimer disease pathology: a bidirectional relationship. Nat Rev Neurol 10(2), 115–119. 10.1038/nrneurol.2013.269

Kandel, E. R., Schwartz, J. H., Jessell, T. M., Siegelbaum, S. A., & Hudspeth, A. J. (2013). Principles of Neural Science (5th ed.). McGraw-Hill Education. 10.1086/670559

Katzman, R., Robert, T., DeTeresa, R., Brown, T., Peter, D., Fuld, P., et al. (1988). Clinical, pathological, and neurochemical changes in dementia: a subgroup with preserved mental status and numerous neocortical plaques. Annals of Neurology,23(2), 138– 144. 10.1002/ana.410230206

Kievit, R. A., Davis, S.W., Mitchell, D.J., Taylor, J.R., Duncan, J., Cam-CAN, & Henderson, R.N.A. (2014). Distinct aspects of frontal lobe structure mediate age-related differences in fluid intelligence and multitasking. Nature Communications, 5, 1–10. 10.1038/ncomms6658

King, J. R., Sitt, J. D., Faugeras, F., Rohaut, B., El Karoui, I., Cohen, L., … & Dehaene, S. (2013). Information sharing in the brain indexes consciousness in noncommunicative patients. Current Biology, 23(19), 1914–1919. 10.1016/j.cub.2013.07.075

Lacaux, C., Strauss, M., Bekinschtein, T. A., & Oudiette, D. (2024). Embracing sleep-onset complexity. Trends in Neurosciences.

Lempel, A., & Ziv, J. (1976). On the complexity of finite sequences. IEEE Transactions on information theory, 22(1), 75–81. 10.1109/tit.1976.1055501

Luppi, A. I., Mediano, P. A., Rosas, F. E., Allanson, J., Pickard, J. D., Williams, G. B., … & Stamatakis, E. A. (2021). Paths to oblivion: common neural mechanisms of anaesthesia and disorders of consciousness. BiorXiv, 2021-02. 10.1101/2021.02.14.431140

Luppi, A. I., & Stamatakis, E. A. (2021). Combining network topology and information theory to construct representative brain networks. Network Neuroscience, 5(1), 96–124. 10.1162/netn_a_00170

Medaglia, J. D., Gu, S., Pasqualetti, F., Ashare, R. L., Lerman, C., Kable, J., & Bassett, D. S. (2016). Cognitive control in the controllable connectome. 10.48550/arXiv.1606.09185

Moran, M., Lynch, C. A., Walsh, C., Coen, R., Coakley, D., & Lawlor, B. A. (2005). Sleep disturbance in mild to moderate Alzheimer’s disease. Sleep medicine, 6(4), 347–352. 10.1016/j.sleep.2004.12.005

Nasreddine, Z. S., Phillips, N. A., Bédirian, V., Charbonneau, S., Whitehead, V., Collin, I., … & Chertkow, H. (2005). The Montreal Cognitive Assessment, MoCA: a brief screening tool for mild cognitive impairment. Journal of the American Geriatrics Society, 53(4), 695–699. 10.1111/j.1532-5415.2005.53221.x

Nelson, H. E., & Willison, J. (1991). National adult reading test (NART). Windsor: Nfer-Nelson. 10.1002/gps.930070713

Nilsson, J. P., Soderstrom, M. Karlsson, A.R. Lekander, M., Akerstedt, T., Lindroth, N.E. & Axelsson, J. (2005). Less effective executive functioning after one night’s sleep deprivation. J. Sleep. Res. 14(1), 1–6. 10.1111/j.1365-2869.2005.00442.x

Pascovich, C., Castro-Zaballa, S., Mediano, P. A., Bor, D., Canales-Johnson, A., Torterolo, P., & Bekinschtein, T. A. (2022). Ketamine and sleep modulate neural complexity dynamics in cats. European Journal of Neuroscience, 55(6), 1584–1600. 10.1111/ejn.15646

Pedroni, A., Bahreini, A., & Langer, N. (2019). Automagic: Standardized preprocessing of big EEG data. NeuroImage, 200, 460–473. 10.1101/460469

Rubinov, M., & Sporns, O. (2010). Complex network measures of brain connectivity: uses and interpretations. Neuroimage, 52(3), 1059–1069. 10.1016/j.neuroimage.2009.10.003

Satz, P. (1993). Brain reserve capacity on symptom onset after brain injury: a formulation and review of evidence for threshold theory. Neuropsychology, 7(3), 273. 10.1037//0894-4105.7.3.273

Sayanng, L. (2009). Sounds downloaded by the time shade. Extracted from: https://freesound.org/people/thetimeshade/downloaded_sounds/

Scarmeas, N., Levy, G., Tang, M.X., Manly, J. & Stern, Y. (2001). Influence of leisure activity on the incidence of Alzheimer’s disease. Neurology, 57(12), 2236–2242. 10.1212/wnl.57.12.2236

Schartner, M., Seth, A., Noirhomme, Q., Boly, M., Bruno, M., Laureys, S., & Barret, A. (2015). Complexity of MultiDimensional Spontaneous EEG Decreases during Propofol Induced General Anaesthesia. PLoS One 10(8). 10.1371/journal.pone.0133532

Schartner, M. M., Carhart-Harris, R. L., Barrett, A. B., Seth, A. K., & Muthukumaraswamy, S. D. (2017). Increased spontaneous MEG signal diversity for psychoactive doses of ketamine, LSD and psilocybin. Scientific reports, 7, 46421. 10.1038/srep46421

Schaefer, A., Kong, R., Gordon, E. M., Laumann, T. O., Zuo, X. N., Holmes, A. J., … & Yeo, B. T. (2018). Local-global parcellation of the human cerebral cortex from intrinsic functional connectivity MRI. Cerebral cortex, 28(9), 3095–3114. 10.1093/cercor/bhx179

Seli, P., Cheyne, J.A., Barton, K.R., & Smilek, D. (2012). Consistency of sustained attention across modalities: comparing visual and auditory version of the SART. Canadian Journal of Experimental Psychology, 66(1), 44–50. 10.1037/a0025111

Shin, H. W., Kim, H. J., Jang, Y. K., You, H. S., Huh, H., Choi, Y. J., … & Hong, J. S. (2020). Monitoring of anesthetic depth and EEG band power using phase lag entropy during propofol anesthesia. BMC anesthesiology, 20(1), 1–10. 10.1186/s12871-020-00964-5

SoundJay (2009). Beep 9. Retrieved from: https://www.soundjay.com/beep-sounds-1.html 10.3998/mpub.10069132.cmp.24

Steffener, J., & Stern, Y. (2012). Exploring the neural basis of cognitive reserve in aging. Biochimica et Biophysica Acta (BBA)-Molecular Basis of Disease, 1822(3), 467–473. 10.1016/j.bbadis.2011.09.012

Stern, Y. (2012). Cognitive reserve in ageing and Alzheimer’s disease. Lancet Neurol, 11(11), 1006–1012. 10.1016/s1474-4422(12)70191-6

Stern, Y., Barnes, C. A., Grady, C., Jones, R. N., & Raz, N. (2019). Brain reserve, cognitive reserve, compensation, and maintenance: operationalization, validity, and mechanisms of cognitive resilience. Neurobiology of Aging, 83, 124–129. 10.1016/j.neurobiolaging.2019.03.022

Tournier, J. D., Smith, R., Raffelt, D., Tabbara, R., Dhollander, T., Pietsch, M., … & Connelly, A. (2019). MRtrix3: A fast, flexible and open software framework for medical image processing and visualisation. Neuroimage, 202, 116137. 10.1016/j.neuroimage.2019.116137

Valenzuela, M. J., Jones, M., Rae, W. W. C., Graham, S., Shnier, R., & Sachdev, P. (2003). Memory training alters hippocampal neurochemistry in healthy elderly. Neuroreport, 14(10), 1333–1337. 10.1097/00001756-200307180-00010

Valenzuela, M. J., & Sachdev, P. (2006). Brain reserve and dementia: a systematic review. Psychological medicine, 36(4), 441–454. 10.1017/s0033291705006264

Veraart, J., Novikov, D. S., Christiaens, D., Ades-Aron, B., Sijbers, J., & Fieremans, E. (2016). Denoising of diffusion MRI using random matrix theory. Neuroimage, 142, 394–406. 10.1016/j.neuroimage.2016.08.016

Whalley, L. J., Deary, I. J., Appleton, C. L., & Starr, J. M. (2004). Cognitive reserve and the neurobiology of cognitive aging. Ageing research reviews, 3(4), 369–382. 10.1016/j.arr.2004.05.001

Winkler, A. M., Ridgway, G. R., Webster, M. A., Smith, S. M., & Nichols, T. E. (2014). Permutation inference for the general linear model. Neuroimage, 92, 381–397. 10.1016/j.neuroimage.2014.01.060

WHO. Fact sheet: dementia. https://www.who.int/news-room/fact-sheets/detail/dementia; 2019, accessed on July 11th, 2020.

Yeh, F. C., Wedeen, V. J., & Tseng, W. Y. I. (2011). Estimation of fiber orientation and spin density distribution by diffusion deconvolution. Neuroimage, 55(3), 1054–1062. 10.1016/j.neuroimage.2010.11.087

Yeo, B. T., Krienen, F. M., Sepulcre, J., Sabuncu, M. R., Lashkari, D., Hollinshead, M., … & Buckner, R. L. (2011). The organization of the human cerebral cortex estimated by intrinsic functional connectivity. Journal of neurophysiology. 10.1152/jn.00338.2011

Yushkevich, P. A., Piven, J., Hazlett, H. C., Smith, R. G., Ho, S., Gee, J. C., & Gerig, G. (2006). User-guided 3D active contour segmentation of an atomical structures: significantly improved efficiency and reliability. Neuroimage, 31(3), 1116–1128. 10.1016/j.neuroimage.2006.01.015

