## Supplementary Material for "Informational Complexity as a Neural Marker of Cognitive Reserve"

[a]Consciousness and Cognition Lab, Department of Psychology, University of Cambridge, Cambridge, UK, [b]Centre for Neuroscience in Education, Department of Psychology, University of Cambridge, Cambridge, UK, [c]Department of Computing, Imperial College London, London, UK, [d]German Center for Neurodegenerative Diseases, Germany, [e]School of Psychology, University of Sussex, Brighton, UK, [f]Institute of Cognitive Neuroscience, University College London, London, UK, [g]Memory Lab, Department of Psychology, University of Cambridge, Cambridge, UK, [h]Department of Psychology, Queen Mary University of London, London, UK

### Introduction

The following supplementary sections provide additional analyses that complement the primary results presented in the main text of 'Informational complexity as a neural marker of cognitive reserve'. We carried out a comprehensive analysis to identify key predictors for sustaining performance under neural strain. In the main text we focused on informational complexity of the EEG signal, as measured by the LZSUM algorithm, as a key predictor. In the supplementary sections, we focused on proxies of cognitive reserve, and structural brain measures.

We computed network measures of white matter connectivity, using the Brain Connectivity Toolbox (Rubinov & Sporns, 2010), to investigate various network characteristics, including characteristic path length, average degree, global efficiency, mean clustering coefficient, mean participation coefficient, small-worldness, network density, and core-periphery structure. For a detailed overview of these measures in the context of cognition, see Rubinov and Sporns (2010). These metrics were calculated for both the overall brain network and seven distinct subnetworks as defined by Yeo et al. (2011), using the Brain Connectivity Toolbox (Rubinov & Sporns, 2010). Additionally, our analysis included structural volumetric MRI data of cortical and medial temporal lobe (MTL) regions. To further contextualise our findings within the framework of cognitive reserve, we included various proxies of cognitive reserve, namely two distinct IQ measures: Cattell (Cattell, 1973) and NART (Nelson & Willison, 1991) along with the discrepancy between them (NART – Cattell), the scores on the Montreal Cognitive Assessment (MoCa) (Nasreddine, 2005), and educational level.

We calculated correlations between the four metrics of behavioural performance differences comparing alert versus drowsy (mean RT, CV RT, commission errors, and omission errors) and: 1) network measures of white matter connectivity, 2) grey matter volume in cortical areas and the MTL, and 3) proxies of cognitive reserve. 4) LZSUM whole brain and 10 regions of interest: (left and right) frontal, central, parietal, temporal, and occipital.

Note that for these performance differences, a larger value indicates greater impairment under drowsy conditions (e.g., slower responses or more errors), so a positive correlation would suggest that worse performance under drowsiness is associated with higher volume, a higher network measure value, a higher LZSUM increase when drowsy, or a higher value on a proxy of cognitive reserve (e.g. IQ, years of education).

### S1 Network measures of white matter connectivity

Correlations were calculated between the 4 performance metrics differences (each for drowsy vs. alert) and network measures of white matter connectivity for both the whole brain and the 7 subnetworks as defined by Yeo et al. (2011) to investigate the potential association between task performance and underlying structural network characteristics. Underlying structural network characteristics can affect the dynamics of functional networks and in this way influence the efficiency of information transfer (Bassett & Sporns, 2017). Various significant correlations and trends were observed, although none survived Bonferroni correction. Positive associations between the mean RT, CV RT, omission errors and the FPC were observed in small-worldness. The core-periphery measure in the VEN showed a consistent positive correlation with CV RT, mean RT, and omission errors. There was also a negative correlation between the mean participation coefficient in the FPC and mean RT difference and a positive correlation between the mean clustering coefficient in both the FPC and DOR and omission errors. Although these correlations did not reach statistical significance after correction for multiple comparisons (Bonferroni and False Discovery Rate), they revealed provisionally interesting patterns laying the groundwork for further investigation into the role of various network characteristics in cognitive reserve. A complete overview of the various correlations and trends between performance metrics and network measures in the different sub-networks is represented in Table S1 and figure S1.

**Table S1 – Correlations between Network Measures and Performance – Uncorrected**

| Performance | Sub-network | Graph theoretic Measure | Correlation |
|---|---|---|---|
| **CV RT** | | | |
| | SOM | Mean clustering coefficient | 0.324† |
| | VEN | Core periphery | 0.365* |
| | FPC | Small-worldness | 0.361* |
| | DMN | Mean participation coefficient | 0.345† |
| **Mean RT** | | | |
| | DOR | Mean clustering coefficient | 0.332† |
| | VEN | Core periphery | 0.432* |
| | FPC | Efficiency | 0.305† |
| | FPC | Mean participation coefficient | -0.385* |
| | FPC | Small-worldness | 0.331† |
| **Omission Errors** | | | |
| | Whole Network | Mean clustering coefficient | 0.321† |
| | Whole Network | Mean participation coefficient | 0.323† |
| | VIS | Mean participation coefficient | 0.374† |
| | DOR | Efficiency | 0.362† |
| | DOR | Mean clustering coefficient | 0.413* |
| | VEN | Core periphery | 0.432* |
| | FPC | Efficiency | 0.376† |
| | FPC | Mean clustering coefficient | 0.358† |
| | FPC | Small-worldness | 0.415* |
| **Commission Errors** | | | |
| | LIM | Characteristic path length | 0.336† |
| | LIM | Core periphery | -0.364† |

**Trend† ($0.05 < p < 0.10$), $p \leq 0.05$ *, $p \leq 0.01$**, $p \leq 0.001$*****

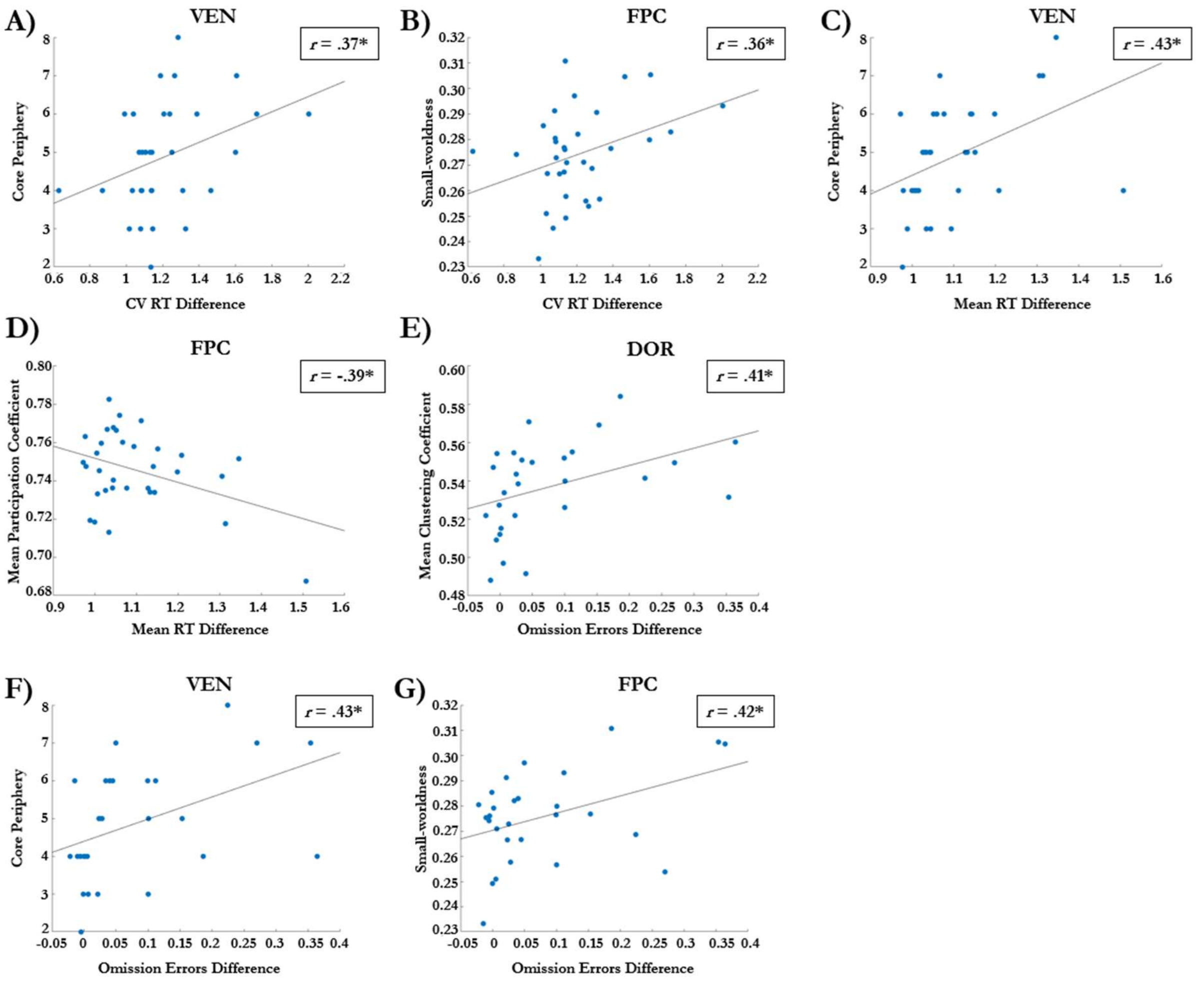


**Figure S1 Scatterplots of associations between network measures of white matter connectivity and performance difference (drowsy vs alert) on various performance metrics.** All panels display scatterplots which illustrate the significant relationship between specific graph-theoretical measures of white matter connectivity (y-axis) and various cognitive performance metrics (x-axis). None of the significant results displayed in the scatterplots survived Bonferroni correction. **A.** Association between core periphery structure in the Ventral Attention (VEN) Network and CV RT difference. **B.** Association between small-worldness in the Frontal Parietal Control (FPC) Network and CV RT difference. **C.** Association between core periphery structure in the Ventral Attention (VEN) Network and mean RT difference. **D.** Association between the mean participation coefficient in the Frontal Parietal Control (FPC) Network and mean RT difference. **E.** Association between the mean clustering coefficient and the Dorsal Attention (DOR) Network and omission errors difference. **F.** Association between core periphery structure in the Ventral Attention (VEN) Network omission errors difference. **G.** Association between small-worldness in the Frontal Parietal Control (FPC) Network and omission errors difference. **($p \leq 0.05$ *), ($p \leq 0.01$**), ($p \leq 0.001$***). Note:** Significance asterisks reflect **uncorrected** p-values (none survived correction).

### S2 Volumetric Analysis

Correlations were calculated between the 4 performance metrics differences (drowsy vs. alert) and cortical and MTL grey matter volume to investigate the potential association between task performance and grey matter volume in key regions (e.g. frontal pole, caudal anterior cingulate) as a potential supporting mechanism in cognitive reserve. Cortical and hippocampal grey matter volume can have a significant effect on cognitive performance and a decrease in grey matter volume is related to healthy aging (Gautam et al., 2011). Exploratory volumetric analyses revealed associations with performance metrics in the Inferior Parietal (L), Lateral Occipital (R), Insula (R), Lingual (L), Parstriangularis (L), Entorhinal (L), Supra Marginal (L), Inferior Parietal (L), and the Paracentral (R). These correlations did not survive correction for multiple comparisons when using both Bonferroni and False Discovery Rate. However, they still reveal potentially interesting patterns that could be further investigated to gain more insight into the potential role of cortical grey matter volume in cognitive reserve. A complete overview of the various associations between performance metrics and grey matter volume is represented in Table S2 and figure S2.

**Table S2. Correlations between Volume and Performance – Uncorrected**

| Performance | Brain Area | Correlation |
|---|---|---|
| **CV RT** | | |
| | Inferior Parietal (left) | -0.42* |
| | Frontal Pole (left) | 0.31† |
| | Insula (left) | -0.31† |
| | Lateral Occipital (right) | 0.44* |
| | Insula (right) | -0.39* |
| **Mean RT** | | |
| | Entorhinal (left) | 0.36† |
| | Inferior Temporal (left) | 0.32† |
| | Lingual (left) | 0.4* |
| | Pars triangularis (left) | -0.35* |
| | Caudal anterior cingulate (right) | 0.31† |
| | Entorhinal (right) | 0.32† |
| | Lingual (right) | 0.31† |
| | Pericalcarine (right) | 0.32† |
| | Precuneus (right) | 0.33† |
| **Commission Errors** | | |
| | Entorhinal (left) | 0.37* |
| | Pars triangularis (left) | -0.39* |
| | Supra Marginal (left) | 0.52** |
| | Inferior Parietal (left) | 0.4* |
| | Paracentral (right) | 0.38* |
| | Supra Marginal (right) | 0.33† |
| | Frontal Pole (right) | 0.33† |
| | Perirhinal cortex | 0.33† |
| **Omission Errors** | | |
| | Caudal anterior cingulate (left) | -0.34† |
| | Superior Frontal (left) | 0.33† |
| | Cuneus (right) | 0.35† |
| | Pericalcarine (right) | 0.35† |

**Trend† (0.05 < p < 0.10), p ≤ 0.05 *, p ≤ 0.01**, p ≤ 0.001*****

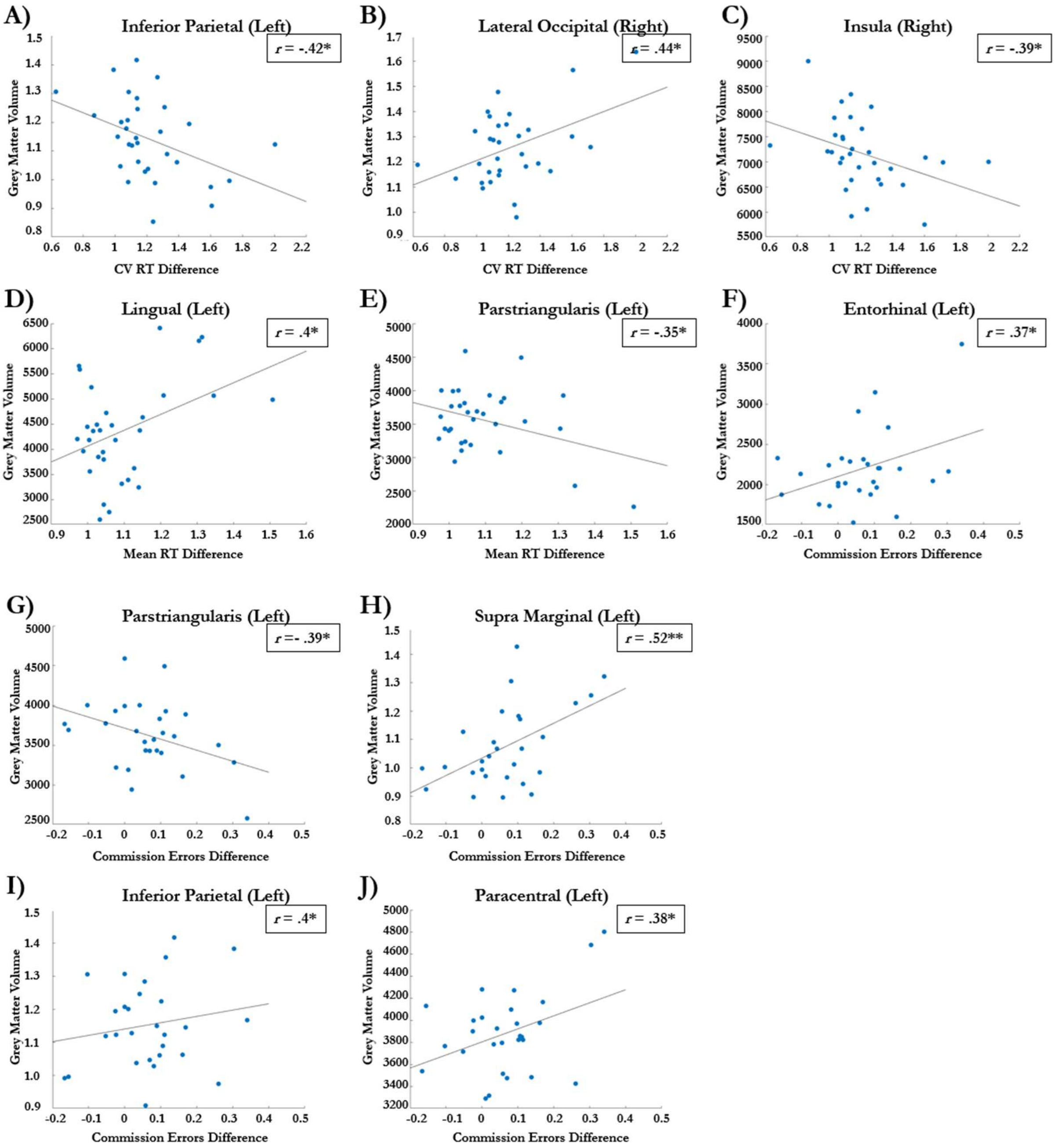


**Figure S2. Scatterplots of associations between volumetric measures and performance difference (drowsy vs alert) on various performance metrics.** All panels display scatterplots which illustrate the significant relationship between grey matter volume of specific brain areas (y-axis) and various cognitive performance metrics (x-axis). None of the significant results displayed in the scatterplots survived Bonferroni correction. **A.** Association between left inferior parietal volume and CV RT difference. **B.** Association between right lateral occipital volume and CV RT difference. **C.** Association between right insula volume and CV RT difference. **D** Association between left lingual volume and mean RT difference. **E.** Association between left pars triangularis volume and mean RT difference. **F.** Association between left entorhinal volume and commission errors difference. **G.** Association between left pars triangularis volume and commission errors difference. **H.** Association between left supra marginal volume and commission errors difference. **I.** Association between left inferior parietal volume and commission errors difference. **J** Association between left paracentral volume and commission errors difference. **($p \leq 0.05$ *), ($p \leq 0.01$**), ($p \leq 0.001$***). Note:** Significance asterisks reflect **uncorrected** p-values (none survived correction).

### S3 Correlational analysis of LZSUM and task performance

We further performed correlations between LZUM of the various ROIs and the 4 metrics of performance. See table S3 for an overview of the uncorrected results. Correlations that survived correction for multiple comparisons are highlighted in bold. See the main text for a further discussion of these results.

**Table S3. LZSUM Correlation Results - Uncorrected**

| Performance | LZSUM | Correlation |
|---|---|---|
| **CV RT** | | |
| | Temporal Right | -0.39* |
| **Mean RT** | | |
| | **Frontal Left** | **-0.53**** |
| | **Frontal Right** | **-0.49**** |
| | **Central Left** | **-0.49**** |
| | Central Right | -0.40* |
| | Temporal Right | -0.35* |
| **Commission Errors** | | |
| | Whole Brain | -0.40* |
| | Frontal Left | -0.43* |
| | Frontal Right | -0.47* |
| | **Central Left** | **-0.56**** |
| | **Central Right** | **-0.60***** |
| | Temporal Right | -0.40* |
| **Omission Errors** | | |
| | Whole Brain | -0.42* |
| | **Frontal Left** | **-0.60**** |
| | **Frontal Right** | **-0.56**** |
| | **Central Left** | **-0.67***** |
| | **Central Right** | **-0.54**** |
| | Temporal Right | -0.47* |

**Trend† (0.05 < p < 0.10), p ≤ 0.05 *, p ≤ 0.01**, p ≤ 0.001*** , correlations that survived Bonferroni are highlighted in bold**

### S4 Proxies of Cognitive Reserve

We explored the relationship between some well-known proxies of cognitive reserve and our measure of cognitive reserve defined as the degree to which task performance was impaired under drowsiness compared to alertness. Specifically, we correlated the 4 metrics of task performance difference (drowsy vs. alert) with age, years of education, fluid intelligence (Cattell), estimation of premorbid IQ (NART), IQ difference (NART – Cattell), and cognitive performance on the MoCa. Importantly, no significant correlations were found between any of the proxies of cognitive reserve and our measure of cognitive reserve.

### Conclusion

We explored correlations between cognitive reserve proxies like IQ and educational level, and performance differences under varying alertness (alert vs. drowsy) conditions. No significant correlations were identified. There could be multiple explanations for the lack of a significant association between these proxies and task performance, such as low statistical power or too much homogeneity in our sample in educational level and IQ (bright old participants), or we may have been measuring something else entirely, such as the ability to compensate for drowsiness on an attentional task. Future studies might benefit from including a broader range of cognitive reserve proxies and expanding the participant profile to include individuals with varying levels of education and cognitive impairment to better assess the thresholds and impacts of educational attainment on cognitive reserve. All these points will be addressed elaborately in the discussion section of the main text.

We further explored correlations between cortical and MTL grey matter volume, network measures of white matter connectivity, and performance differences (alert vs. drowsy). Some interesting significant results and trends were observed, but none of these survived Bonferroni correction. Despite the lack of statistically significant results in volumetric and network analyses, these preliminary trends observed that indicate further research could elucidate the role of brain structure and function in cognitive reserve. Our discussion in the main text (see Discussion section) elaborates further on the implications of these findings in the context of the main results.

### References


Bassett, D. S., & Sporns, O. (2017). Network neuroscience. *Nature neuroscience*, *20*(3), 353-364. https://doi.org/10.1038/nn.4502

Cattell, R. B. (1973). *Culture-fair intelligence test*. Institute for Personality and Ability Testing. https://doi.org/10.1037/t14354-000

Gautam, P., Cherbuin, N., Sachdev, P. S., Wen, W., & Anstey, K. J. (2011). Relationships between cognitive function and frontal grey matter volumes and thickness in middle aged and early old-aged adults: The PATH Through Life Study. *Neuroimage*, *55*(3), 845-855. https://doi.org/10.1016/j.neuroimage.2011.01.015

Nasreddine, Z. S., Phillips, N. A., Bédirian, V., Charbonneau, S., Whitehead, V., Collin, I., ... & Chertkow, H. (2005). The Montreal Cognitive Assessment, MoCA: a brief screening tool for mild cognitive impairment. *Journal of the American Geriatrics Society*, *53*(4), 695-699. https://doi.org/10.1111/j.1532-5415.2005.53221.x

Nelson, H. E., & Willison, J. (1991). *National adult reading test (NART)*. Windsor: Nfer-Nelson. https://doi.org/10.1002/gps.930070713

Rubinov, M., & Sporns, O. (2010). Complex network measures of brain connectivity: uses and interpretations. *Neuroimage*, *52*(3), 1059-1069. https://doi.org/10.1016/j.neuroimage.2009.10.003

Yeo, B. T., Krienen, F. M., Sepulcre, J., Sabuncu, M. R., Lashkari, D., Hollinshead, M., ... & Buckner, R. L. (2011). The organization of the human cerebral cortex estimated by intrinsic functional connectivity. *Journal of neurophysiology*. https://doi.org/10.1152/jn.00338.2011
